# Single-cell perturbation prediction near experimental reproducibility with GENESIS

**DOI:** 10.64898/2026.10.30.763081

**Authors:** Riccardo Conci, Davide D’Ascenzo, Sebastiano Cultrera di Montesano, Srivatsan Raghavan, Peter S. Winter, Eliezer M. Van Allen, Lorin Crawford, William Lotter

## Abstract

Single-cell perturbation atlases now measure cellular responses across millions of cells, but the number of possible combinations of biological contexts, perturbations, doses and other experimental variables make exhaustive measurements infeasible. Here, we present GENESIS (Generative ENErgy-based Single-cell Intervention Simulator), a generative framework that predicts single-cell response distributions directly from the covariates defining an experimental condition. GENESIS uses a lightweight neural network trained with the energy score, a strictly proper scoring rule for distributions. We introduce an experiment-calibrated evaluation framework in which prediction error is interpreted relative to the reproducibility of repeated measurements of the same condition. Across the Tahoe-100M small-molecule atlas and a large PBMC cytokine atlas, GENESIS significantly outperforms existing generative models and statistical baselines and approaches experimental reproducibility on independently repeated conditions. Importantly, GENESIS remains accurate when trained on only a fraction of the available experimental conditions or cells, and generated profiles recover experimentally observed context-dependent transcriptional programs. At comparable training-cell budgets, GENESIS achieves more accurate predictions when measurements span more experimental conditions rather than profiling fewer conditions more deeply. Together, these findings suggest a complementary strategy for perturbation studies: broader, lower-depth measurements can expand experimental coverage, while GENESIS augments sparsely sampled populations and predicts responses in unmeasured conditions.

## Introduction

Cells remodel their transcriptomes in response to genetic, chemical and environmental perturbations. Single-cell perturbation atlases now measure these responses at unprecedented scale: Tahoe-100M profiles more than 100 million cells across approximately 55,000 drug–cell-line–dose conditions ^1^, while the peripheral blood mononuclear cell (PBMC) cytokine atlas profiles approximately 10 million cells across diverse cytokine perturbations ^2^. Yet even datasets of this scale sample only a small fraction of the possible experimental space, which spans factors such as biological context (such as cell lines or donors), perturbation, dose, timing and other experimental variables. This vast combinatorial space makes exhaustive measurement infeasible. Computational models could instead use measured conditions to predict outcomes across the much larger space of unobserved experimental settings, but this strategy requires both data-efficient learning and a meaningful standard for judging generated predictions.

Several generative approaches have been developed to predict single-cell perturbation responses. Most formulate the problem as learning a mapping from a measured population of control cells to a perturbed population, using set-based transformers ^3^, variational autoencoders ^4^, optimal transport ^5^, flow matching ^6^ or diffusion models ^7^. Yet this mapping is not directly observed experimentally: because single-cell sequencing is destructive, control and perturbed populations consist of different cells rather than paired measurements before and after treatment. Evaluating these models presents a further difficulty ^8–10^: although new methods are routinely benchmarked against held-out perturbation responses, it remains unclear how close we are to predicting responses as reliably as they can be measured. Even when the same cell line is exposed to the same drug and dose on separate experimental plates, the measured responses can differ owing to finite sampling and variation between experimental runs. Without a reference for this experimental reproducibility, improvements over existing methods do not establish how close predictions come to the agreement achievable between independent experimental repeats.

Here, we introduce GENESIS (Generative ENErgy-based Single-cell Intervention Simulator), which models cellular response distributions directly from the covariates defining an experimental condition. In the datasets studied here, these specify the cell line or donor and cell type, the drug or cytokine applied, its dose where applicable, and the experimental batch. A lightweight neural network uses these covariates to generate individual cell profiles without requiring measured control cells at inference. The model is trained using the energy score, a strictly proper scoring rule that enables learning distributions directly from individual observations, without requiring paired cells or fixed-size population targets. The framework accommodates different sets of experimental covariates and incomplete coverage of their possible combinations.

We evaluate GENESIS on Tahoe-100M and the PBMC cytokine atlas, demonstrating improved prediction of held-out experimental conditions relative to existing generative models and statistical baselines. To contextualize performance, we use independent experimental replicates in Tahoe-100M to establish a reproducibility benchmark, which GENESIS approaches. We further show that GENESIS maintains strong performance with substantially reduced training data and that generated transcriptomes capture biologically meaningful, context-dependent responses. Together, these findings establish a framework for predicting cellular responses across subsampled experimental spaces and inform potential strategies for designing more efficient perturbation studies.

### GENESIS generates perturbation responses from experimental covariates

We model a single cell’s transcriptome as a draw from a distribution conditioned on the covariates specifying its experimental condition. For ease of reference, we refer to these covariates in three groups: biological context (e.g., cell line), perturbation (e.g., drug, dose), and experimental replication context (e.g., plate identifier), which collectively define the condition. Denoting a condition as *c* and the cells’ expression as *Y*, GENESIS learns the conditional distribution *P*(*Y* | *c*). Each covariate is represented by a learned embedding, which maps the value of the covariate to a numerical representation. The embeddings defining a condition are concatenated with random noise and passed through a multilayer perceptron to generate a single-cell gene-expression profile (Methods). Repeated noise draws generate populations of arbitrary size (Fig. 1a).

**Fig. 1.**
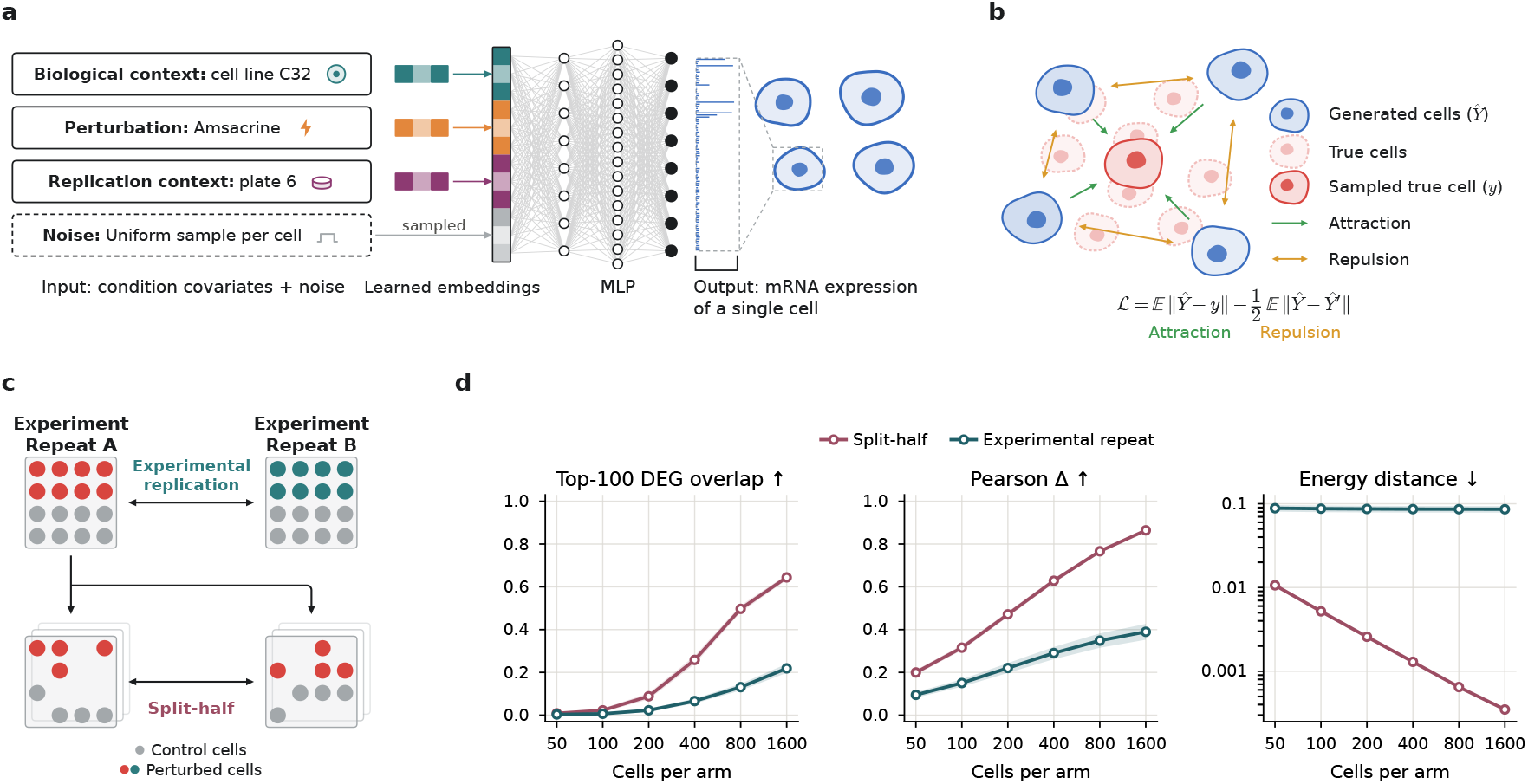
GENESIS generates single-cell perturbation responses from experimental covariates, with prediction performance benchmarked against experimental reproducibility. **a**, Overview of GENESIS. Covariate values describing biological, perturbation and replication context are each represented by a learned embedding; the embeddings are concatenated with a noise vector drawn independently for every cell and passed through a multilayer perceptron that outputs a predicted expression profile of one cell. Repeated noise draws generate a population of any size. **b**, The energy-score objective. The attraction term encourages generated cells to resemble the observed cell *y*, while the repulsion term encourages diversity among generated cells, preventing collapse to a single mean profile and allowing the model to capture cell-to-cell variability. **c**, Two references for experimental agreement. An experimental repeat compares one condition assayed in two independent experiments; a split-half repeat compares two disjoint random halves of the cells from one experiment, capturing cell-to-cell variability and finite sampling but not batch effects. **d**, Experimental and split-half reproducibility as a function of cell count. We analyzed the one hundred most deeply profiled Tahoe-100M conditions measured on two plates, each with at least 3,200 perturbed and 3,200 control cells per plate. At each cell count *n*, we sampled *n* perturbed and *n* control cells per repeat, drawing experimental repeats from separate plates and split-half repeats from non-overlapping subsets of the same plate. Agreement was measured by direction-matched overlap among the top 100 differentially expressed genes, Pearson *Δ*, and energy distance. Means across conditions are plotted after averaging 20 random draws per condition at each cell count; bands indicate the standard error of the mean across conditions. Split-half reproducibility improves more dramatically with cell count, whereas experimental reproducibility remains limited by experimental variation.

This formulation is particularly natural for single-cell perturbation experiments. Because single-cell sequencing is destructive, control and perturbed cells are unpaired samples from condition-level distributions: the pre-perturbation state of an individual assayed cell is never observed. GENESIS therefore requires neither measured control cells at inference nor an assumed correspondence between individual control and perturbed cells. Instead, a control measurement (i.e., the null perturbation) is simply represented using a dedicated value for the perturbation covariate.

GENESIS is trained to recover the full conditional distribution using the energy score ^11^ as a loss function (Methods). The energy score can be computed from a single observed cell of a condition. It balances two requirements: generated transcriptomes should lie close to the observed transcriptome but also differ from one another, which encourages the model to capture variation across sampled cells during training (Fig. 1b). More fundamentally, the energy score is a strictly proper scoring rule: averaged over observations, the true distribution is its unique optimum. This framework thus enables learning heterogeneous condition-level responses directly from individual cells, without paired observations or fixed-size population targets. We additionally include a variogram term ^12^ in the loss function that encourages the model to reproduce the average magnitude of expression differences between pairs of genes within cells, helping capture gene–gene dependence and improving convergence in practice (Methods).

### Experimental reproducibility provides a reference for model performance

Various metrics have been proposed to measure agreement between predicted and measured transcriptomes, including direction-matched differential-expression gene overlap ^3^, control-relative Pearson correlation ^13,14^ and energy distance ^15^. While these metrics each measure useful aspects of concordance, they are difficult to interpret without a reference for what level of agreement is experimentally achievable. We therefore anchor predictive performance to **experimental reproducibility**: how closely an independently repeated experiment reproduces the original measurement (Fig. 1c, Methods). When the same condition is assayed in different experimental batches, such as on different plates in Tahoe-100M, disagreement between the two measurements captures both finite sampling and variation introduced by repeating the experiment ^16,17^. We refer to a prediction as reaching experimental reproducibility when its agreement with the measured response is comparable to that between independent experimental repeats. Because independent repeats are available for only a subset of conditions, we additionally introduce **split-half reproducibility** for conditions measured only once. Cells from a single experiment are randomly divided into two equally sized populations and compared, with the partition redrawn 20 times and results averaged across draws (Fig. 1c, Methods). Split-half reproducibility captures finite-sampling variability without the additional variation introduced by repeating the experiment, hence it provides an overly-optimistic but still informative reference for scenarios lacking experimental reproducibility data.

Fig. 1d compares the experimental and split-half reproducibility benchmarks in Tahoe-100M as a function of the number of cells used in the calculation. Averaged over the one hundred most deeply profiled Tahoe-100M conditions measured on two plates, both benchmarks show relatively high experimental variability, but split-half reproducibility improves more sharply with cell count. Additional cells therefore reduce finite-sampling variability but do not eliminate variation introduced by repeating the experiment. Differential-expression analysis, a routine readout of perturbation experiments, is strongly affected: even at 1,600 cells per arm, two plates share on average only about 20 of their top 100 differentially expressed genes, compared with more than 60 for two halves of the same plate. These findings highlight the importance of experimental reproducibility as a reference for model performance.

### GENESIS performance approaches experimental reproducibility

We first evaluated GENESIS on its ability to predict perturbation responses in held-out biological contexts (i.e., cell lines for Tahoe-100M and donors for PBMC), following the train/test splits and selected gene panels introduced in prior work ^3,7^. For each cell line or donor selected for evaluation, controls and a minority of randomly selected perturbations are observed during training, while the majority of perturbations are reserved for testing. The model must therefore infer unobserved responses using perturbation measurements from other biological contexts and the limited training measurements available for the same cell line or donor (Methods, Extended Data Fig. 1a,c). We benchmarked GENESIS against PerturbDiff ^7^, STATE ^3^ and CellFlow ^6^, together with linear, mean-based and nearest-neighbor baselines (Methods, Supplementary Methods). We scored predictions with three metrics that capture complementary aspects of a response: direction-matched top-100 differentially expressed gene (DEG) overlap ^3^, which asks whether predictions recover the genes a perturbation significantly changes and the direction of their effect; control-relative Pearson correlation (Pearson *Δ*)^13,14^, which measures agreement across genes between the predicted and observed mean expression changes relative to matched controls; and energy distance ^15^, which compares full single-cell distributions, including differences beyond their means (Methods, Extended data Fig. 2).

We first assessed how closely GENESIS performance approaches the reproducibility of the experiments themselves. Averaged across independently repeated Tahoe-100M conditions, GEN-ESIS performance reached experimental reproducibility in Pearson *Δ* (GENESIS: 0.430 [95% CI 0.381-0.482]; experimental reproducibility: 0.432 [0.378-0.491]; *p* = 0.56) and energy distance (GENESIS: 0.158 [0.147-0.170]; experimental: 0.154 [0.143-0.166]; *p* = 0.36), and fell only 3.4% short in the top-100 DEG overlap (GENESIS: 0.289 [0.263-0.317]; experimental: 0.299 [0.271-0.330]; *p* = 8 × 10^−4^) (Fig. 2a, Methods, Supplementary Table 1). All other models and baselines differed significantly from experimental reproducibility across all three metrics. Experimental reproducibility itself varied substantially across conditions. When plate pairs were ranked by agreement between experimental repeats, GENESIS tracked this variation most closely (Fig. 2b).

**Fig. 2.**
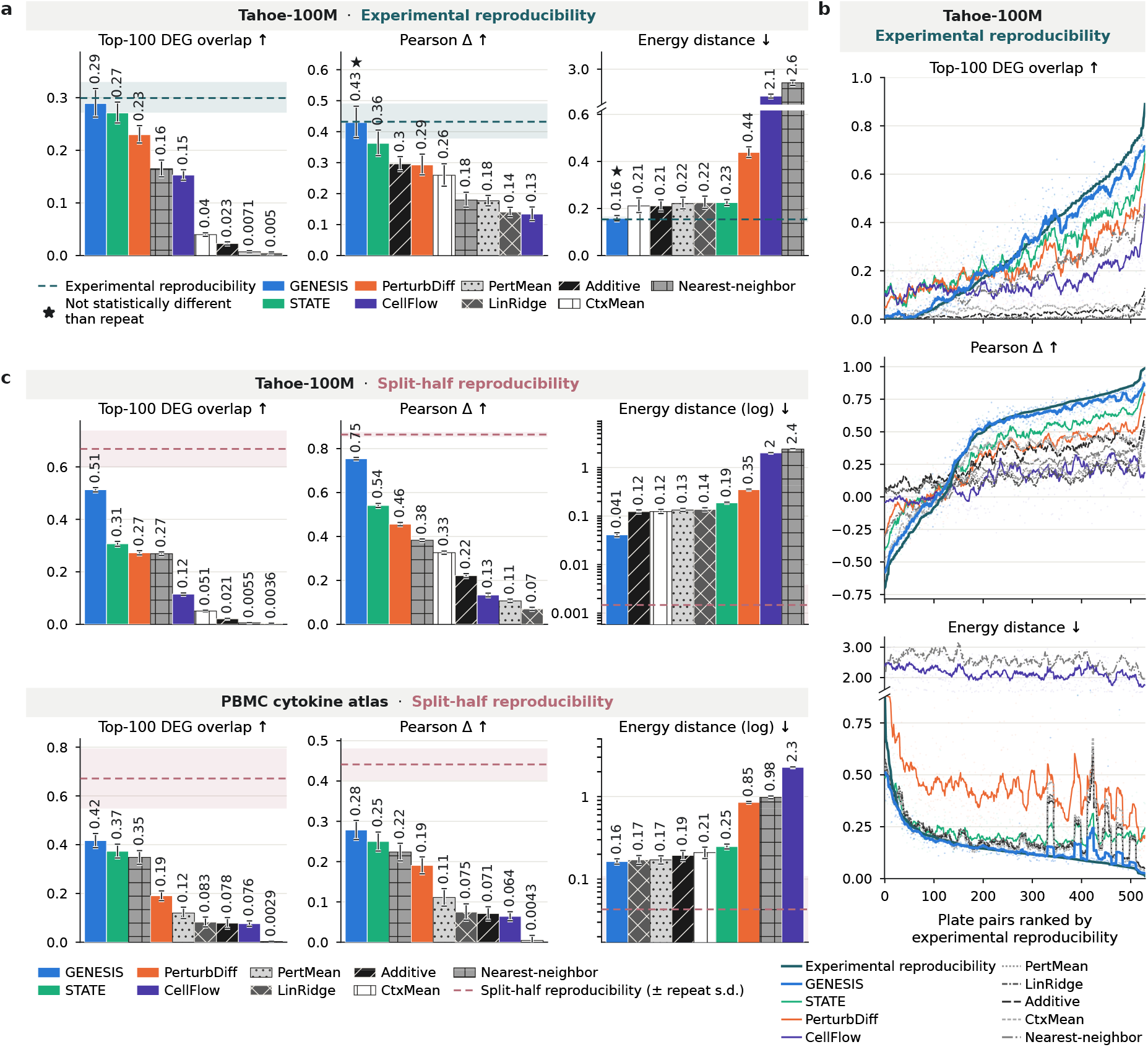
GENESIS predicts cellular responses in unseen conditions at a level approaching experimental reproducibility. **a**, Tahoe-100M held-out cell lines, scored against experimental reproducibility. For 530 pairs of plates measuring the same condition, each model generates one plate from that plate’s controls and is scored against the other, real plate; experimental reproducibility (dashed line; band, 95% CI) scores the two real plates against each other. ⋆, not statistically different from experimental reproducibility (*P* ≥ 0.05) in both the mean or the win rate (the share of plate pairs on which the model is closer to the truth than the other real plate is; 50% for a second replicate). Both tests are paired, two-sided and clustered by combination. Every other model differed from experimental reproducibility on every metric (*P* < 0.001). **b**, The same 530 plate pairs ranked by experimental reproducibility on each metric. Model curves are computed as a centred running mean (averaging the 13 points each direction). GENESIS tracks experimental reproducibility most closely of all models (Pearson *r* between model and experimental reproducibility across the 530 per-pair scores: 0.95, 0.98 and 0.66 for top-100 DEG overlap, Pearson *Δ* and energy distance), ahead of STATE (0.80, 0.92, 0.47), PerturbDiff (0.69, 0.88, 0.46) and CellFlow (0.52, 0.56, 0.18). **c**, Held-out contexts scored against split-half reproducibility in Tahoe-100M held-out cell lines (4,211 conditions; 4,064 for DEG overlap) and PBMC cytokine atlas held-out donors (4,268 conditions; 2,395 for DEG overlap). Split-half reproducibility scores two random halves of a condition’s cells against each other, repeated over 20 random splits (dashed line, mean; band, ± s.d. across the 20 splits, averaged over conditions). Energy distance is on a log axis. Bars, mean; error bars, 95% cluster-bootstrap CI (10,000 resamples; clusters are cell line × compound in Tahoe-100M and donor × cytokine in PBMC). ↑/↓, higher/lower is better.

Experimental replicates cover only a subset of the held-out conditions in Tahoe-100M (*N* = 530) and are absent from the PBMC cytokine atlas, where each donor was assayed on its own plate, so plate and donor cannot be separated (Extended Data Fig. 3). We therefore next evaluated all held-out conditions (*N* = 4211 on Tahoe-100M, *N* = 4268 on PBMC cytokine atlas), using split-half reproducibility to contextualize performance. GENESIS showed state-of-the-art performance on both datasets (Fig. 2c). In Tahoe-100M, direction-matched top-100 DEG overlap increased from an average of 30.7 genes [95% CI 29.7-31.6] for the next-best method (STATE) to 51.3 [50.5-52.1] for

GENESIS, a 67% increase; in the PBMC cytokine atlas, from 37.3 [34.5-40.2] to 41.7 [38.8-44.6], a 12% increase. The split-half reproducibility values are 66.8 and 67.1 for Tahoe-100M and PBMC, respectively. GENESIS also achieved the highest Pearson *Δ* and the lowest energy distance on both datasets (Fig. 2c).

We further evaluated a complementary perturbation-generalization setting. Whereas the previous setting withholds most perturbation responses from selected cell lines or donors for testing, here we withhold most cell lines or donors from selected perturbations for testing. Specifically, in Tahoe-100M, test perturbations were observed in only 1, 5, 15 or 20 training cell lines, while in the PBMC atlas, test perturbations were observed in only 1, 3 or 5 training donors (Methods; Extended Data Fig. 1b,d). Across 4,023 held-out conditions in Tahoe-100M and 3,456 in the PBMC cytokine atlas, GENESIS again outperformed all competing methods on all three metrics (Extended Data Fig. 4).

### GENESIS maintains strong performance with reduced training data

We next investigated how predictive performance depends on the amount of training data. To test biological-context generalization, we retrained GENESIS on reduced versions of the Tahoe-100M dataset. Here, we used either fewer experimental conditions or fewer cells per condition, while matching the control and perturbed-cell budgets and keeping the original test set fixed (Methods). Remarkably, using only ∼5% of the available perturbed training cells (4.6 million of 89.5 million), GENESIS outperformed all competing generative models and statistical baselines trained on the full dataset across all three metrics (Fig. 3a, Supplementary Table 2). Relative to the strongest full-data competitors, GENESIS increased top-100 DEG overlap by approximately 50% and Pearson *Δ* by 30% (STATE), while reducing energy distance by approximately 60% (context-mean baseline). Thus, GENESIS achieved state-of-the-art performance with 20-fold fewer perturbed-cell measurements. At these reduced data budgets, the allocation of measurements also mattered. Sampling fewer cells across more conditions improved performance relative to deeply profiling fewer conditions, particularly at ∼5% of the training data, whereas the difference narrowed at ∼25% (Fig. 3a). Taking this to the extreme, we trained GENESIS using all controls and at most 25 cells per perturbed condition, corresponding to approximately 1.3 million perturbed cells across the training set. Augmenting these conditions with GENESIS-generated cells to their original sizes recovered both DEG overlap and Pearson *Δ*, bringing them close to experimental reproducibility (Extended Data Fig. 5). This suggests that GENESIS can recover responses from very shallow measurements by leveraging information shared across the broader experimental space.

**Fig. 3.**
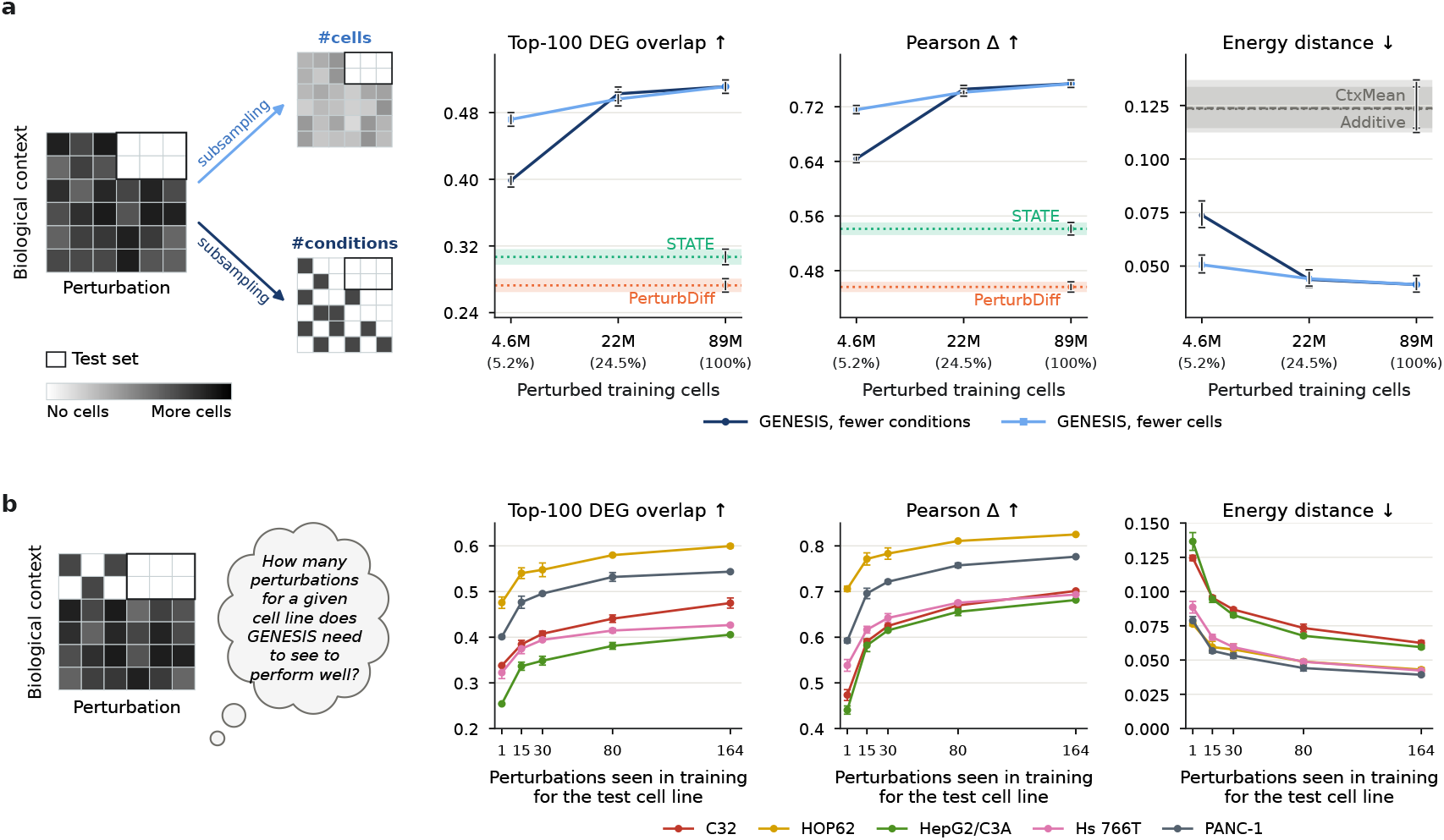
GENESIS retains state-of-the-art performance with sparse experimental coverage. **a**, Effect of reducing the amount of training data on Tahoe-100M. GENESIS is trained on data subsampled either by reducing the number of cells per condition or by reducing the number of conditions, with the total number of training cells matched across the two settings and the test set held fixed. When subsampling, we first targeted 5% and 25% of conditions, which resulted in slightly different percentages of total cells given variable numbers of experimental cells per condition (Methods). Points show mean performance across three runs with different model-training and subsampling seeds; whiskers indicate 95% cluster-bootstrap confidence intervals over conditions, clustered by cell line × compound. For each performance metric, results are also shown for the two highest-performing existing models or baselines. **b**, Effect of training overlap for individual biological contexts. For each held-out test cell line, the number of perturbations observed for that cell line during training is progressively increased while the evaluated perturbations remain held out. Perturbations included in training are selected at random across three independent seeds. Performance improves rapidly with the first observed perturbations and shows diminishing gains as overlap increases, with little variation across random seeds.

Finally, we asked how many perturbations must be measured in a given cell line before GENESIS can accurately predict its responses to others. For each of the five held-out Tahoe-100M cell lines, we progressively increased the number of perturbations observed during training while keeping the test set fixed (Methods). Observing just 15 perturbations, fewer than 10% of the 164 available training compounds, captured approximately half the performance gain obtained with all 164, with diminishing improvements thereafter (Fig. 3b). Performing a similar analysis under the perturbation-generalization setting showed a similar pattern: observing a perturbation in as few as five cell lines already enabled strong predictions of its responses in other, held-out cell lines (Extended Data Fig. 6).

### Generated profiles recapitulate context-dependent biological programs

The value of generated single-cell profiles ultimately depends on whether they support the same biological interpretations as measured cells. We therefore asked whether GENESIS could reproduce transcriptional responses at the level of signaling pathways, a common readout of perturbation experiments. We used PROGENy ^18^, which infers pathway activity from experimentally derived transcriptional footprints, providing a measure of downstream signaling responses rather than the expression of pathway components themselves. We evaluated all 14 pathways across 55 compounds in the five held-out Tahoe-100M cell lines, quantifying responses as PROGENy-weighted mean log_2_ fold changes relative to matched controls. For this analysis, we expanded the utilized gene panel to encompass those used in PROGENy-defined pathways and trained GENESIS on only 5% of the available training cells (“fewer cells” configuration in Fig. 3a; Methods). Generated profiles closely reproduced the measured patterns of pathway induction and suppression (Fig. 4). Quantitatively, across conditions measured on independent plates, experimental repeats showed an average Pearson correlation of 0.455 [95% CI 0.415-0.499] across the 14 pathway responses. GENESIS achieved a comparable average correlation of 0.402 [0.370-0.439] with the measured responses, outperforming all other models and statistical baselines (Extended Data Fig. 7).

**Fig. 4.**
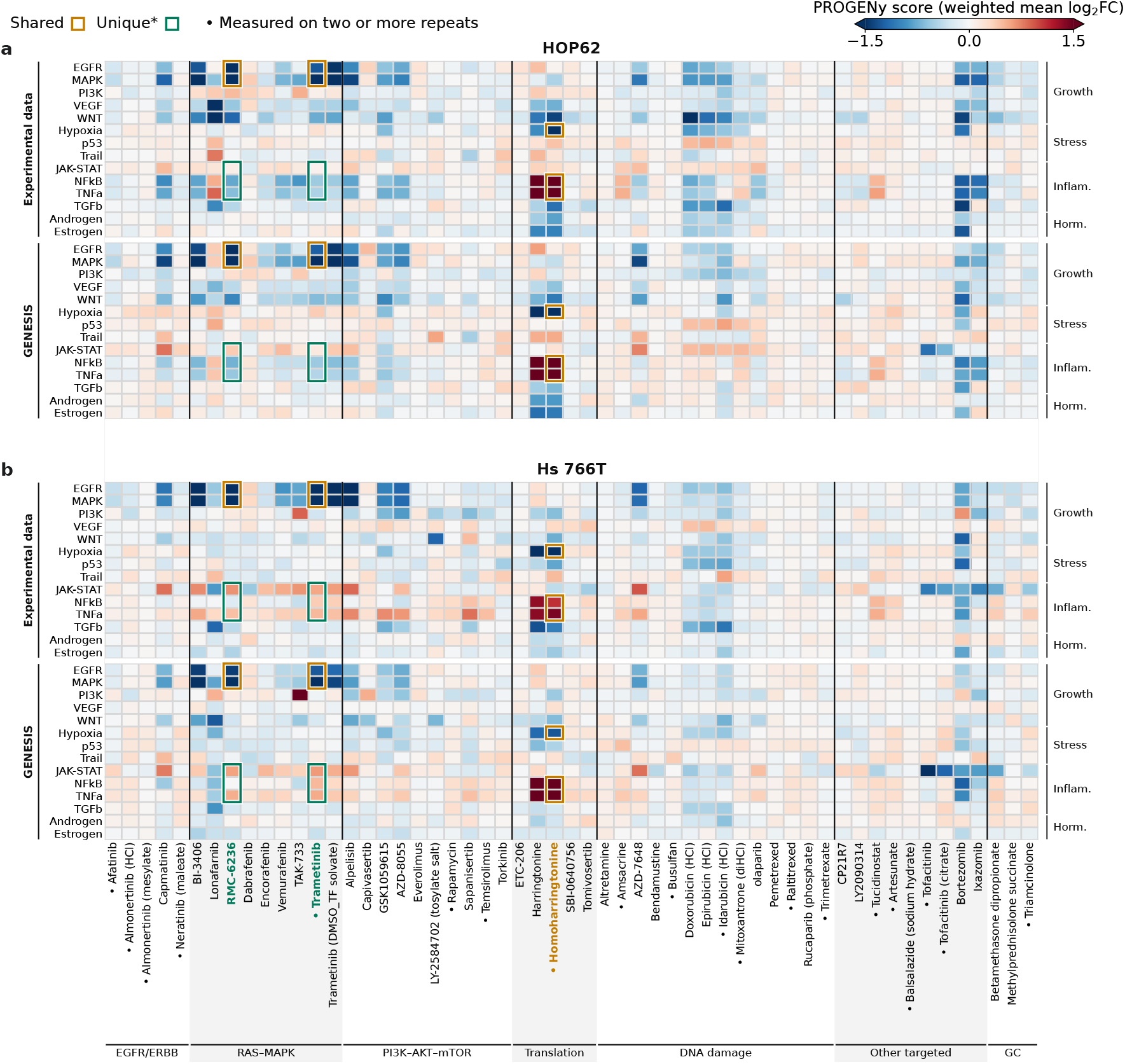
GENESIS recovers shared and line-specific pathway responses in held-out KRAS-mutant cell lines. Measured and GENESIS-predicted PROGENy pathway responses in HOP62 (lung adenocarcinoma, KRAS G12C) and Hs 766T (pancreatic carcinoma, KRAS Q61H) across 55 compounds at 5 µM. Both lines were held out in the Tahoe-100M biological-context setting; no line-compound combination shown was observed during training. GENESIS was trained on 5% of training cells using a 3,072-gene panel. For each line, measured responses are shown above predictions. Rows show the 14 PROGENy pathways grouped by function; columns show compounds grouped by class and alphabetically within each class (GC for Glucocorticoids). • marks compounds measured on two or more plates (20/55). Colors show pathway-module log_2_ fold changes relative to same-plate DMSO controls, calculated as PROGENy-weighted means over the top 100 footprint genes detected in at least 10% of control cells. Before scoring, gene-level log_2_ fold changes were depth-normalized for both measured responses and predictions (Methods). Colors are clipped at ±1.5. Boxes highlight selected compound-pathway pairs discussed in the main text meeting criteria for shared responses (amber: both lines change in the same direction by at least 0.5 in log_2_ fold change) or line-specific responses (green: between-line difference of at least 0.5 in log_2_ fold change). Corresponding entries are boxed in measured and predicted panels; compound labels match the box color, with green used when both categories apply.

We next asked whether GENESIS could distinguish responses shared across biological contexts from those specific to individual cell lines. This distinction is particularly important in cancer, where the same treatment can produce different downstream effects depending on genetic background. We examined two KRAS-mutant cell lines, HOP62 lung cancer cells and Hs 766T pancreatic cancer cells, which exhibited both shared and distinct responses (Fig. 4). The MEK inhibitor trametinib and the RAS(ON) inhibitor RMC-6236 (daraxonrasib) ^19^ both suppressed EGFR- and MAPK-associated transcriptional activity in the two lines, and GENESIS reproduced these shared responses (Fig. 4). Yet their inflammatory responses diverged: both treatments increased JAK-STAT and TNFα pathway scores in Hs 766T, while suppressing them in HOP62. GENESIS captured these opposing responses, despite not observing these compound-cell-line combinations during training (Fig. 4).

## Discussion

GENESIS combines a conditional generator with a distributional training objective to predict single-cell perturbation responses for unseen combinations of experimental covariates. Across both Tahoe-100M and the PBMC cytokine atlas, it outperforms existing generative and statistical approaches and, when independent repeats are available, achieves performance comparable to experimental reproducibility. Because GENESIS conditions directly on recorded experimental covariates, it can also be readily adapted to different experimental designs.

Beyond predictive performance, our pathway analysis suggests that GENESIS recovers context-dependent transcriptional programs. The captured differences between HOP62 and Hs 766T cell lines may be particularly relevant to cancer therapy, where compensatory signaling can influence responses to targeted inhibitors. The suppression of TNFα-associated transcription in HOP62 is consistent with the established oncogenic role of KRAS in promoting inflammatory signaling ^20–22^. Conversely, increased JAK-STAT activity in Hs 766T is consistent with reports of compensatory STAT3 activation following MAPK inhibition, a potential resistance mechanism that has motivated combined MEK and JAK/STAT3 inhibition in pancreatic cancer ^23,24^ and is now under further clinical investigation ^25^. The response to RMC-6236 also parallels interferon-associated signaling reported in pancreatic tumors following prolonged RAS inhibition, which has been linked to adaptive cell-state changes and therapeutic resistance ^26^. By recovering these contrasting responses alongside shared MAPK suppression, GENESIS preserves biologically meaningful differences between cellular contexts that may influence the effects of targeted therapies.

Our findings also have potential implications for experimental design. Once finite-sampling noise falls below between-experiment variation, measuring additional conditions may be more informative than sequencing more cells per condition. Consistent with this principle, GENESIS retains strong performance even with substantially reduced training data, especially when reducing cells per condition rather than the number of conditions.

Important areas of future work remain. GENESIS generalizes compositionally: an unseen combination of covariates can be predicted only when each covariate value has appeared elsewhere during training. It therefore cannot yet predict responses to a completely unseen compound or to a new cell line or donor. Incorporating external information about perturbations and biological contexts, such as the chemical structure of compounds or cell-line genotypes, could extend the framework beyond this setting. More generally, many variables that shape cellular responses, including duration of treatment exposure, culture medium and features of the cellular environment, are not often recorded in perturbation datasets and therefore cannot be explicitly modeled by GENESIS. Recording these factors would enable their incorporation as additional covariates, potentially transforming hidden sources of experimental variation into predictable dimensions of cellular response.

Together, our findings establish GENESIS as an accurate and data efficient framework for predicting single-cell perturbation responses across unmeasured experimental conditions. By approaching experimental reproducibility while retaining biologically meaningful variation, GENESIS demon-strates the potential of generative models to extend the reach of perturbation atlases beyond directly measured conditions.

## Methods

### Datasets and data processing

#### Tahoe-100M

The Tahoe-100M dataset ^1^ comprises 100,648,790 cells from 50 cancer cell lines exposed to 379 compounds at three doses (0.05, 0.5 and 5 µM) across 14 plates, each containing all 50 lines. A condition was defined by the combination of cell line, compound, dose and plate, yielding 65,243 perturbed conditions. Dimethyl sulfoxide (DMSO) vehicle controls were matched to perturbed conditions by cell line and plate.

#### PBMC cytokine atlas

The Parse Biosciences peripheral blood mononuclear cell (PBMC) cytokine atlas ^2^ comprises 9,697,974 cells from 12 donors, each assayed on a 96-well plate containing 90 cytokine stimulations and six phosphate-buffered saline (PBS) control wells. Cells were annotated by the dataset authors into 18 cell types. A condition was defined by donor, cytokine, cell type and well; the well identifier was retained to distinguish the six independent PBS controls from each donor. Controls were matched by donor and cell type. Because each donor was assayed on a separate plate, donor and plate effects are confounded and the dataset contains no independent between-plate repeat of the same biological condition (Extended Figure 3).

#### Gene selection and preprocessing

For the main benchmarking analyses, we used the same 2,000 highly variable genes (HVGs) selected in prior work ^3,7^, enabling direct comparison with existing methods. GENESIS was trained on log1p-transformed raw UMI counts of the selected genes, without library-size normalization. Training cells were streamed using scDataset (v0.3.0) ^27^ and sampled uniformly without replacement within each epoch, without reweighting experimental conditions (Supplementary Table 3).

### Perturbation prediction tasks

We considered two complementary prediction tasks. In biological-context generalization, most perturbation responses were withheld from selected cell lines or donors, while their controls and a minority of responses remained available for training. In perturbation generalization, selected compounds or cytokines were observed in only a limited number of biological contexts during training, with their responses withheld in the remaining contexts.

#### Biological-context generalization

For Tahoe-100M, we used the context-generalization split of STATE ^3^ to enable direct comparison with prior work. This split uses five cell lines for evaluation, with test responses drawn from 268 compounds: 199 are withheld at all three doses, whereas 69 are withheld only at 0.05 µM and 0.5 µM, with their 5 µM responses remaining in training. A further 19 compounds are reserved for validation, which we used for hyperparameter tuning. Each evaluation cell line retains its controls and responses to 164 training compounds. In total, the split comprises 4,211 test conditions (Extended Data Fig. 1a). When comparing model performance with experimental reproducibility, we restricted the evaluation to test conditions measured independently on multiple plates, excluding compounds observed at any dose in the corresponding evaluation cell line during training. This yielded 500 cell line-compound-dose combinations measured on two or three plates with at least 50 cells per plate, resulting in a total of 530 independent plate pairs.

For the PBMC cytokine atlas, we likewise used the donor-generalization split of STATE ^3^. This split uses four donors (1, 4, 9 and 12) for evaluation. For each evaluation donor, responses to 61 cytokines are withheld for testing, four cytokines are reserved for validation, and responses to the remaining 25 cytokines are retained for training. The remaining eight donors provide training responses across the cytokine panel. This yields 244 held-out donor-cytokine combinations, evaluated separately by cell type. PBS controls from all donors remained in training (Extended Data Fig. 1c).

#### Perturbation generalization

To evaluate generalization across perturbations, we constructed a complementary split with training and test sets comparable in size to those of the biological-context split, but with limited observations of each test perturbation across cell lines. For Tahoe-100M, we withheld 40 randomly selected compounds at all three doses in 30 test cell lines, yielding 4,035 test conditions. For each compound, training responses were retained in 1, 5, 15 or 20 of the remaining 20 cell lines, randomly selected for each compound, with ten compounds assigned to each level. All other compounds were observed in all 50 cell lines (Extended Data Fig. 1b). The experimental reproducibility benchmark for this task comprised 403 held-out cell line–compound–dose combinations measured on two plates with at least 50 cells per plate. Because all three doses were withheld in each corresponding test cell line, no response to these compounds had been observed in those cell lines during training.

For the PBMC cytokine atlas, we followed a similar design. Thirty randomly selected cytokines were withheld in seven donors, giving 210 test donor–cytokine combinations. Each cytokine was observed during training in 1, 3 or all 5 of the remaining donors, while the other 60 cytokines remained available across all donors (Extended Data Fig. 1d).

### GENESIS

#### Conditional generator

For each experimental condition *c*, GENESIS generates individual cell profiles by combining learned representations of the experimental covariates with random noise. The covariates specify the experimental condition, while independent noise draws allow the model to generate different cells from the corresponding distribution. For Tahoe-100M, the covariates consisted of a cell line (e.g., C32), a compound (e.g., amsacrine), a dose and a plate. For the PBMC cytokine atlas, the covariates consisted of a donor, a cell-type, a cytokine and a well.

Each covariate value is mapped to a learned *d*-dimensional embedding, with *d* = 256 and coordinates initialized independently from *N*(0, 1/*d*). Thus, the initialization standard deviation is 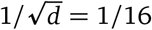. These embeddings are concatenated with a *d*-dimensional noise vector whose coordinates are drawn independently from 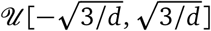. Since a uniform distribution on [−*a, a*] has variance *a*^2^/3, these bounds give the noise coordinates variance 1/*d*, matching the embedding coordinates at initialization. Consequently, each input block has unit expected squared norm, so the embeddings and noise enter the network on the same scale at initialization.

A multilayer perceptron with hidden widths 4,096 and 8,192 and GELU activations generates an expression profile, with a rectified linear output giving non-negative log(1 + count) values. No observed cells are required as inputs. Controls are represented by a dedicated perturbation value, and independent noise draws generate cells from the same condition.

#### Energy score

For an observed cell *y* from condition *c*, the energy score ^11^ is given by

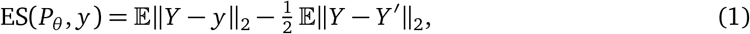

where *Y* and *Y* ^′^ are independent cells generated by GENESIS for condition *c*, that is, independent draws from its conditional distribution *P*_θ_ (· | *c*), where θ denotes the model’s trainable parameters. Distances are computed across panel genes in log(1 + count) space. The first term pulls generated cells towards the observation; the second rewards spread among them. The energy score is a strictly proper scoring rule: for distributions with finite first moments, its expected value is uniquely minimized by the true distribution. In training, both expectations were replaced by averages over *m* cells ŷ_1_, …, ŷ_*m*_ generated for condition *c*,

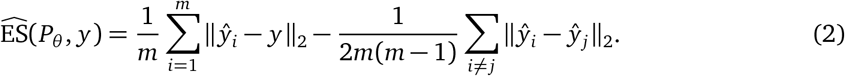

Excluding self-pairs from the generated-sample distance term yields an unbiased estimator of the energy score ^28^. GENESIS uses *m* = 256 generated samples per observation by default, with sensitivity analysis presented in Extended Data Fig. 14a.

#### Variogram term

Although the energy score is strictly proper, accurately learning gene-gene dependencies can be challenging with finite training data. We therefore added a variogram score to emphasize these dependencies by comparing within-cell expression differences across gene pairs ^12^. For genes *g* and *h* of a cell *y*, let *v*_*gh*_(*y*) = | *y*_*g*_ − *y*_*h*_|^1/2^, with | *y*_*g*_ − *y*_*h*_| floored at 10^−4^. The variogram score

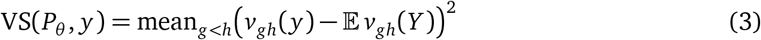

is proper, so ES + λ VS remains strictly proper.

To obtain an unbiased estimate of the variogram score, we must account for the finite number of generated cells. Estimating E*v*_*gh*_(*Y*) by averaging over the *m* generated cells and then squaring the residual would add Var[*v*_*gh*_(*Y*)]/*m* to the expected loss, introducing an artificial penalty on variability in the generated distribution. Instead, we split the generated cells into two independent groups, obtaining separate estimates 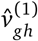 and 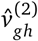, and multiply the corresponding residuals,

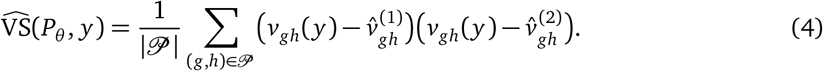

This yields an unbiased estimator because the two estimates are independent. At each training step, *P* contained 6,000 gene pairs sampled using random perfect matchings of the panel genes, and 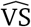 was computed on half of the observed cells in each minibatch. The full training objective combines the energy score and variogram estimators:

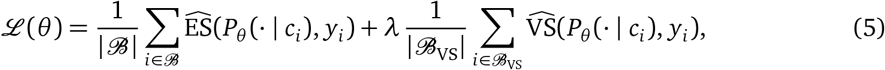

where *ℬ* is the minibatch of observed cells and *ℬ*_VS_ ⊆ *ℬ* contains half of those cells. We used λ = 0.5, chosen empirically from the validation set. Sensitivity analysis of this variable is shown in Extended Data Fig. 14b).

#### Training configuration

The Tahoe-100M GENESIS benchmark model was trained for five epochs (256,205 optimization steps) using AdamW (β_1_ = 0.9, β_2_ = 0.95; weight decay 10^−4^ on weight matrices only; no gradient clipping), with batches of 1,792 observed cells, 256 generated cells per observed cell and bfloat16 mixed precision. The learning rate increased linearly from zero to 9 × 10^−4^ over the first 5% of optimization steps, with a minimum of 100 and a maximum of 1,000 warm-up steps. It was then reduced to 0.1 and 0.01 of its peak value at approximately 20% and 40% of training, respectively, followed by a linear decay to 0.1 of its current value over the final 15% of steps. We evaluated the final checkpoint without validation-based model selection. Default training settings are reported in Supplementary Table 3.

On a single NVIDIA H100 NVL GPU, GENESIS processed approximately 2,500-3,000 observed cells per second. Training on the Tahoe-100M biological-context split (91.8 million cells) took approximately 43 hours, while training on the PBMC split (7.3 million cells) took approximately 4 hours.

For the training-overlap experiments (Figure 3b), which required repeated training runs, we used a *fast-training configuration* with 96 generated cells per observed cell instead of 256 and a larger batch size of 2,048. The architecture and training objective were unchanged; training duration was adjusted as described in Supplementary Methods.

### Comparator models and baselines

#### Deep learning methods

We compared GENESIS with three generative models that predict perturbed cells from observed control cells: State ^3^, a transformer-based model; PerturbDiff ^7^, a diffusion model; and CellFlow ^6^, a flow-matching model. For biological-context generalization, we used released checkpoints for STATE and PerturbDiff on Tahoe-100M and for PerturbDiff on the PBMC atlas. We retrained STATE on PBMCs and trained CellFlow on both datasets. For perturbation generalization, all three methods were retrained from scratch. Training used the published implementations and recommended configurations.

All methods were evaluated on the same test conditions and gene panels, generating the same number of cells per condition. For all methods, we followed the authors’ training and inference procedures. In particular, for Tahoe100M, input covariates for all models included the plate. For the PBMC cytokine atlas, STATE (through cell-type-matched control cells) and PerturbDiff (through a cell-type covariate) were conditioned on PBMC cell type. Following the authors’ procedure for CellFlow on this dataset, we conditioned the model on donor and cytokine and assigned cell-type labels to generated cells by nearest-neighbor label transfer in CellFlow’s 100-dimensional PCA space. Specifically, each generated cell inherited the fine cell-type label of its nearest training cell, using up to 1,000 training cells per condition as reference ^6^.

#### Statistical baselines

We compared GENESIS with five baselines fitted only on training conditions, weighting each condition equally. The *context mean* predicts the average expression across training perturbations in the same cell line and plate. The *perturbation mean* adds the average response to the target perturbation across training contexts to the target context’s control mean. The *additive model* combines the control mean with an overall mean response and separate context and perturbation effects, without an interaction term. *Linear ridge regression* ^14^ estimates the perturbation response in a 100-dimensional gene space derived from training pseudobulks, with regularization strength 0.1. *Nearest-neighbor* selects the training context most similar to the target context across their shared training perturbations and copies its measured cells under the target perturbation on the same plate. Similarity was measured by cosine distance between responses projected onto 100 principal components. Full equations and fitting procedures for all baselines are provided in Supplementary Methods.

### Evaluation

#### Evaluated populations

For each comparison, every method generated as many cells as the observed population it is compared with. Observed and generated populations were compared with the observed controls of the same cell line and plate (Tahoe-100M) or donor and cell type (PBMCs). Conditions with fewer than four perturbed or control cells were excluded.

#### Integer counts

Generated values were converted to integer counts before scoring. Outputs in log(1 + count) were rounded to the integer nearest in log(1 + count), that is, to ⌊*v*⌋ or ⌈*v*⌉, whichever minimizes | log(1 + *k*) −log(1 + *v*)|. Outputs in counts (the baselines, and comparators after their normalization was inverted) were rounded to the nearest integer; negative values were set to zero. Observed counts are integers already.

#### Depth-scaled log space

For each cell, let **x** denote the vector of raw UMI counts over the selected gene panel. We transformed these counts as

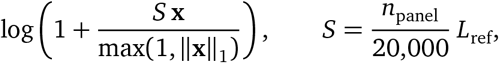

where *n*_panel_ = 2,000 and *L*_ref_ is the median full-transcriptome library size of the dataset (1,829 for Tahoe-100M and 3,320 for PBMCs). *S* approximates the expected panel library at median sequencing depth: 183 and 332, respectively. Scaling every cell to *S* removes differences in sequencing depth between plates and methods, so metrics computed in this space compare expression composition.

#### Evaluation metrics

We compared an observed population *T* and a generated population 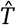 of the same condition with three metrics. Two of them are relative to controls; for these, the matched control cells were split at random into disjoint halves, *C* for the observed and *Ĉ* for the generated population, so that the two responses do not share a control estimate.

##### DEG overlap

In each population, every gene detected in at least 10% of all matched control cells was tested against the population’s control half with a two-sided, tie-corrected Mann-Whitney U test on counts ^29,30^, a test that performs well for single-cell differential expression ^31,32^, and *P* values were adjusted across genes by the Benjamini-Hochberg procedure ^33^. A gene was differentially expressed if *q* < 10^−3^ and the absolute log_2_ fold change of mean counts was at least 0.1 (pseudocount 10^−9^). The fold-change threshold follows the default of Seuratv5^34^, and the stricter *q* threshold guards against the inflated significance of cell-level tests ^35^. DEGs were identified separately for the observed and predicted responses: the former by comparing measured perturbed cells with matched controls, and the latter by comparing generated cells with matched controls. After ranking the DEGs in each population by *q*, let *D*_*k*_ and 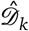 denote the top *k* = min(100, *n*_obs_) genes in the observed and predicted lists, respectively, where *n*_obs_ is the number of observed DEGs.

We define

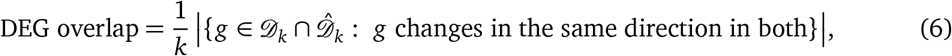

which is undefined when *n*_obs_ = 0.

##### Control-relative Pearson correlation

With 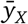 the mean expression of population *X* in the depth-scaled log space, the observed and predicted responses are

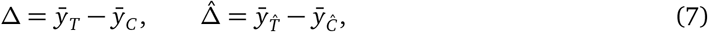

and *r*_*Δ*_ is their Pearson correlation across panel genes.

##### Energy distance

For observed cells *x*_1_, …, *x*_*n*_ and generated cells *y*_1_, …, *y*_*m*_ in the depth-scaled log space, each is subsampled at random to at most 4096 cells for computational efficiency. Most conditions contained fewer than 4096 cells, so this cap affected only larger populations and limited the quadratic computational cost of pairwise distance calculations. We then computed energy distance as

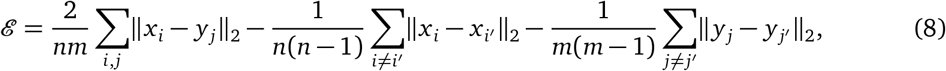

which is the unbiased estimator of 2 E∥*X* − *Y* ∥_2_ − E∥*X* − *X* ^′^∥_2_ − E∥*Y* − *Y* ^′^∥_2_. Controls do not enter this metric.

#### Experimental reproducibility

For each condition measured on two or more Tahoe-100M plates, we evaluated all available plate pairs separately. Let *A* and *B* denote the observed cell populations from the same condition measured in two plates, and *Â* and 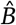 the corresponding model-generated populations, each matched in size to its observed counterpart. For each evaluation metric *m*, we define

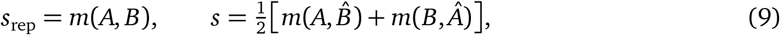

where *s*_rep_ measures agreement between independent experimental repeats, while *s* averages prediction performance in both cross-plate directions. Thus, predictions for one plate are evaluated against measurements from the other, with control-relative metrics using each plate’s matched controls. The benchmark comprises 530 plate pairs (Methods, “Perturbation prediction tasks”).

The performance of each method with respect to experimental reproducibility was summarized by the ratio of means, *R* = Σ *s*/ Σ *s*_rep_, and its “win rate” which is the fraction of cross-plate predictions that outperform the repeat (higher DEG overlap or *r*_*Δ*_, lower energy distance; ties count as half). Because plate pairs from the same combination share plates, both were tested against no difference (*R* = 1; win rate 50%) by two-sided *t*-tests with standard errors clustered by combination and *G* − 1 degrees of freedom ^36,37^. *R* was tested on the log scale, with its standard error computed using the delta method ^38^.

#### Split-half reproducibility

The perturbed cells and matched controls of each test condition were split at random into disjoint halves, *T*_1_, *T*_2_ and *C*_1_, *C*_2_. The reference score is *m*(*T*_1_, *T*_2_), with *C*_1_ and *C*_2_ for control-relative metrics, averaged over 20 random partitions. A method is scored on the first partition by replacing *T*_2_ with an equal-sized generated population. All eligible test conditions were included, each plate’s population as a separate condition, and each metric was averaged over the conditions for which it was defined for every method and the reference.

### Data efficiency and scaling

#### Fewer conditions or fewer cells

To ask whether a fixed experimental budget is better spent measuring more conditions or more cells per condition, we reduced the Tahoe-100M biological-context training set in two ways while keeping the test set unchanged. In the fewer-conditions setting, we retained all cells from either 5% or 25% of the 52,885 perturbed training conditions, defined by cell line, compound and dose with plates pooled. Conditions were selected using hierarchical balanced stratified sampling (see next paragraph). In the fewer-cells setting, we retained the same number of perturbed cells, sampled uniformly across all training conditions. All control cells were retained in both settings. The two training sets therefore contained the same numbers of control and perturbed cells and differed only in how broadly the perturbed measurements were distributed across conditions. Reduced datasets were trained for 5/ *f* epochs, where *f* is the retained fraction of conditions, ensuring at least as many optimizer steps as for the full training set.

#### Hierarchical balanced stratified sampling

When subsampling conditions, we wanted to reduce the size of the training set without preferentially retaining the most densely measured parts of the dataset. We therefore selected conditions so that coverage remained approximately balanced across cell lines, compounds and doses, while also encouraging each compound to be represented across diverse tissue types. The sampler first assigns inclusion weights to all measured cell line, compound and dose combinations so that their expected representation is balanced along each of these three axes. For each compound and dose, these weights determine how many cell lines are retained. Cell lines are then sampled without replacement, with lower probability given to tissues already represented for that compound. This procedure is repeated until the target fraction of conditions is reached. The full algorithm is given in Supplementary Methods.

#### Recovery from shallow measurements

We trained GENESIS using all control cells and at most 12 perturbed cells per training condition, pooled across plates, yielding approximately 1.3 million perturbed cells (Extended Data Fig. 5a, b). To assess recovery of training-condition responses, we generated populations matching their original sample sizes and evaluated 3,941 combinations measured on plates 6 and 14 at 5 µM. We compared these predictions with the shallow measurements alone, the same measurements replicated to the held-out sample size, and GENESIS trained on all cells. Measured-cell baselines were scored against cells excluded from training; model predictions were scored against all measured cells. Generalization was evaluated separately on the unchanged biological-context test set. Both analyses used cross-plate experimental reproducibility as the reference.

#### Reducing the total cell budget

To assess data efficiency when control measurements are also limited, we uniformly retained 1%, 5%, 25% or 100% of all training cells, including both perturbed and control cells. Models were evaluated on the unchanged biological-context test set using split-half reproducibility (Extended Data Fig. 5c).

#### Training overlap

To measure how prediction improves as a biological context is observed under more perturbations, we varied the number of training compounds available for each of the five Tahoe-100M test cell lines while keeping the evaluation set fixed. Each line retained responses to *k* ∈ {0, 15, 30, 80, 164} of its 164 training compounds, at all available doses, with *k* = 0 containing controls only. Compound sets were nested, so each higher-overlap level contained all compounds from the preceding levels. The same 199 compounds withheld from these lines at all doses were used for evaluation at every level. To separate the effect of training overlap from differences between cell lines, we used a Latin-square design across five runs, such that every cell line was evaluated once at every overlap level.

We repeated the complete design for three independently sampled compound orderings, giving 15 runs in total.

For the perturbation-generalization analysis (Extended Data Fig. 6a), we similarly varied the number of biological contexts in which a test compound was observed during training. The four overlap levels were rotated across four groups of ten held-out compounds and repeated for three independent selections of training cell lines, giving 12 runs. All Tahoe-100M overlap experiments used the same uniformly subsampled set of 11.3 million training cells and the lighter training configuration.

For the PBMC data-efficiency analysis (Extended Data Fig. 6b), models were trained on either 5% or 25% of cells sampled uniformly from the full training set, or on 25% of donor-cytokine conditions selected using the same balanced sampling procedure, without the tissue-balancing term.

### Biological program analysis

#### Tahoe-100M PROGENy programs

##### Pathway footprints and gene panel

We scored the 14 signaling pathways of PROGENy ^18^. Each pathway is defined by a footprint of genes whose expression changes consistently when the pathway is perturbed, and each gene carries a signed weight. For each pathway we took the 100 genes with the smallest *P* values in the human model, the package default, as distributed through OmniPath ^39^. Tahoe-100M measures 1,287 of the resulting 1,297 genes. Only 221 of these are in the default 2,000-gene panel used for model development, so we trained and evaluated on the union of the two, 3,066 genes. Six randomly drawn genes padded the panel to a multiple of 64 for computational efficiency and were excluded from every analysis. Before any analysis, we grouped the pathways by the signals that drive them: growth factor (EGFR, MAPK, PI3K, VEGF, WNT), stress (hypoxia, p53, TRAIL), inflammation (JAK-STAT, NF-κB, TNFα, TGFβ), and hormonal and developmental signaling (androgen, estrogen). For this analysis, GENESIS was retrained on the expanded gene panel using the same training data and configuration as the 5% fewer-cells model in Fig. 3a.

##### Compounds

In the five held-out cell lines of the biological-context split (C32, HepG2/C3A, HOP62, Hs 766T and PANC-1), 199 compounds were withheld at every dose, so training included none of their compound-cell-line combinations. We analyzed, at 5 µM, every one of them that the Tahoe-100M annotation ^1^ assigns to one of 13 mechanism classes expected to alter the signaling that PROGENy reads out: inhibitors of EGFR/ERBB, RAS, RAF, MEK, PI3K/AKT, mTOR, JAK/STAT, GSK3, the proteasome, HDAC, DNA synthesis or repair, or protein synthesis, and glucocorticoid receptor agonists. This gave 55 compounds. The classes were fixed before any response was examined. Twenty compounds were measured on two or more 5 µM plates and 35 on one. Each compound was on the same plates in every line, with at least 50 perturbed cells per plate.

##### Pseudobulk transcriptional responses

For each compound–cell line pair at 5 µM, we calculated the mean raw count of each gene across perturbed cells and across DMSO-treated control cells from the same plate. This pseudobulk aggregation reduced the influence of single-cell sparsity, while comparison with plate-matched controls limited the contribution of plate-specific technical variation.

Genes were included in the analysis if they were detected in at least 10% of the corresponding control cells. For compounds measured on multiple plates, control detection fractions were averaged across plates with equal weighting. This criterion was determined exclusively from measured control cells and applied identically to measured and predicted responses.

For each eligible gene *g*, the pseudobulk log-fold change was calculated as

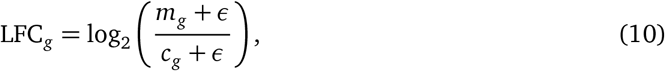

where *m*_*g*_ and *c*_*g*_ denote the mean counts in perturbed and plate-matched control cells, respectively, and ε = 10^−9^ ensured a defined ratio without appreciably altering non-zero expression estimates. For compounds measured on multiple plates, we summed the plate-specific perturbed means and control means before calculating the ratio. Thus, plates contributed equally to the aggregation irrespective of their cell counts.

#### Adjustment for library depth

To account for differences in library depth between perturbed and control cells, we calculated the total transcript count across the whole transcriptome for each measured cell. We then adjusted each gene-level log-fold change by subtracting the log-ratio of mean library depths:

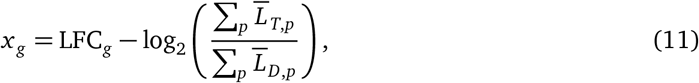

where 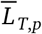 and 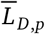 denote the mean total transcript counts in perturbed and DMSO-treated cells on plate *p*, respectively. The sums include all plates on which the compound was measured and reduce to a single term for compounds measured on one plate. This aggregation matched that used to calculate gene-level fold changes.

Apart from the negligible pseudocount, the adjusted response corresponds to the log-fold change in each gene’s fraction of captured transcripts. Library depth was estimated from the whole transcriptome rather than the selected gene panel, avoiding normalization to the aggregate expression of the panel itself.

Because GENESIS generated counts only for the selected panel, whole-transcriptome library depths were unavailable for predicted cells. We therefore applied the library-depth adjustment from each measured condition to its corresponding predicted response. Subtracting the same adjustment from measured and predicted log-fold changes preserved their gene-level differences.

#### Pathway response scores

We summarized the adjusted gene-level responses using the PROGENy footprints defined above. For each pathway, we retained the selected footprint genes represented in the expanded panel and passing the control-detection threshold, without re-ranking genes after restriction to the panel. The pathway response score was calculated as

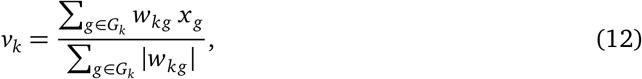

where *G*_*k*_ is the set of eligible footprint genes for pathway *k, w*_*kg*_ is the corresponding PROGENy weight and *x*_*g*_ is the library-depth-adjusted log-fold change. The signed weights encode the direction and magnitude of each gene’s contribution to the pathway footprint. Normalization by the sum of absolute weights retained the log-fold-change scale and accounted for differences in the total weight of eligible genes across pathways and conditions. Measured and predicted scores were calculated using the same eligible genes and weights.

#### Shared and line-specific responses

To illustrate shared and context-dependent responses, we examined HOP62 lung cancer cells and Hs766T pancreatic cancer cells, which harbor KRAS G12C and Q61H mutations, respectively ^40,41^. These two held-out cell lines provided distinct biological contexts in which to examine responses to inhibition of RAS-MAPK signaling. We focused on the MEK inhibitor trametinib and the RAS(ON) inhibitor RMC-6236, highlighting examples of shared pathway suppression and opposing inflammatory pathway responses (Fig. 4).

#### Quantitative evaluation of pathway responses

We evaluated pathway-level agreement using the conditions and cross-plate comparisons described above (Methods: Experimental reproducibility; Extended Data Fig. 7). GENESIS and STATE were trained on the same 5% of training cells using the expanded 3,072-gene panel and compared with the additive and nearest-neighbor baselines. Pathway responses were calculated separately for each plate as described above, retaining footprint genes detected in at least 10% of control cells on both plates. Nearest-neighbor responses used the source line’s controls, with genes additionally required to pass detection in those controls.

We assessed Pearson correlation along three axes: *across pathways*, over the 14 pathway scores for each compound-dose condition and cell line; *across compounds*, over compound–dose conditions for each pathway and cell line; and *across cell lines*, over the five held-out lines for each compound-dose condition and pathway. Correlations were averaged over the remaining dimensions, giving each cell line equal weight where applicable. Model correlations were averaged over both cross-plate directions, and multiple plate-pair estimates were averaged within each condition.

Confidence intervals at the 95% percentile were calculated from 10,000 cluster-bootstrap resamples, using identical resamples across methods and recomputing each statistic. For the across-pathway and across-compound analyses, clusters comprised cell line-compound combinations, retaining all doses together. For the across-cell-line analysis, clusters comprised compounds, retaining all cell lines and doses together.

Finally, to assess whether the recovery of biological programs is robust to a different gene-set collection and scoring method, we complemented the PROGENy analysis with preranked GSEA ^42^ of MSigDB Hallmark gene sets ^43^, comparing predicted enrichment scores with measured responses and experimental reproducibility (Supplementary Methods; Extended Data Fig. 13).

## Data availability

The datasets used in this work were obtained from Huggingface for Tahoe-100M (https://huggingface.co/datasets/tahoebio/Tahoe-100M) and from the author’s website for the Parse PBMC cytokine atlas (https://www.parsebiosciences.com/datasets/10-million-human-pbmcs-in-a-single-experiment/).

## Code availability

The source code for GENESIS is available under the MIT licence via GitHub at https://github.com/rickconci/GENESIS.

## Acknowledgements

S.C.M. was supported by the Eric and Wendy Schmidt Center at the Broad Institute of MIT and Harvard. This research was supported in part by a David & Lucile Packard Fellowship for Science and Engineering (L.C.), the National Library of Medicine (R01LM014775; W.L.) and the Ellison Foundation (E.M.V.A. and W.L.). S.R. acknowledges funding support from NCI K08 CA260442. Any opinions, findings, and conclusions or recommendations expressed in this material are those of the author(s) and do not necessarily reflect the views of any of the funders.

## Author contributions

R.C., D.D. and S.C.M. developed GENESIS. R.C., D.D., S.C.M. and W.L. designed the study and wrote the first draft of the manuscript. R.C. performed the computational experiments and analyses. S.R., P.S.W., E.M.V.A. and L.C. provided scientific feedback and revised the manuscript. All authors reviewed and approved the final manuscript.

## Supplementary Methods

### Training optimization steps

Training on the full Tahoe-100M biological-context split for five epochs required 256,205 optimizer steps. On the reduced dataset retaining 25% of perturbed cells and all controls (approximately 24.2 million cells), validation loss stopped improving after approximately 150,000 steps. We used this plateau to set a shorter training budget and derive a scaling rule for other dataset sizes.

Specifically, we fitted a power law through the full-data budget and the validation-based budget for the 25%-cells dataset:

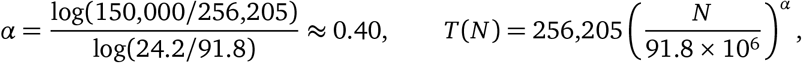

where *N* is the total number of training cells, including controls. PBMC models trained for five epochs (20,425 steps), taking approximately four hours on one NVIDIA H100 NVL.

#### Fast-training configuration

The 27 repeated runs for the training-overlap experiments (Fig. 3b) and related data-efficiency analyses used 96 generated cells per observation and batches of 2,048 observed cells. The architecture, objective and other optimizer settings followed the default configuration.

### Statistical baselines

We compared GENESIS with five statistical baselines (perturbation mean, context mean, linear ridge, additive and nearest neighbor), each fitted on the training conditions and gene panel of the corresponding analysis (the 2,000 panel genes unless stated otherwise). We index biological context by *b*, perturbation by *p*, and experimental replicate by *r*, where *r* in Tahoe-100M identifies a plate. Let *Y*_*bpr*_ and *Ŷ*_*bpr*_ denote the true and predicted cell-expression distributions, and let 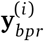 and 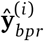 denote individual observed and predicted cells, as counts on the panel genes. Following previous works ^3,14^, we normalize and log-transform each cell’s counts:

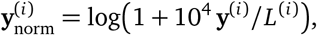

where *L*^(*i*)^ is the cell’s full-transcriptome library size, defined as its total count over all measured genes. We write ***µ***_*bpr*_ for the mean of 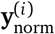 over a condition’s cells and 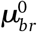 for the same mean over its matched control cells. The observed transcriptomic response is

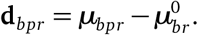

In Tahoe-100M, biological context denotes a cell line, perturbation denotes a compound–dose pair, and controls (DMSO) are matched by cell line and plate. In the PBMC cytokine atlas, biological context denotes a donor-cell-type combination and perturbation denotes a cytokine. Each donor occupies its own plate, so *r* is determined by the donor. Each donor-cytokine condition occupies a single well, and its controls pool cells of the corresponding cell type from the donor’s six PBS wells.

#### Generating populations from pseudobulk vectors

To turn a predicted perturbed pseudobulk into a cell population, we subtract the observed control pseudobulk and add the resulting difference to individual control cells. This shifts the control population toward the predicted perturbed expression while incorporating variation from real cells, allowing evaluation using energy distance and DEG overlap. In normalized log-expression space, each sampled control cell is shifted as

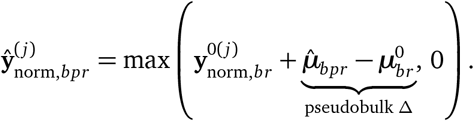

#### Context mean

The context mean predicts the average expression observed across training perturbations in the target biological context. It captures the context’s typical perturbed expression without distinguishing between perturbations. Mathematically,

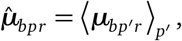

where the average runs over the non-control training perturbations measured in context *b* on plate *r*. If no such conditions are available on the target plate, we average over all non-control training conditions of *b* across plates.

#### Perturbation mean

The perturbation mean assumes that a perturbation induces a shared expression change across biological contexts. It averages the matched-control responses of training conditions containing that perturbation and adds the resulting response to the target context’s control mean. Mathematically,

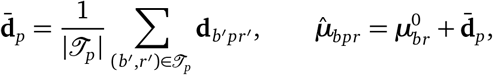

where,*J*_*p*_ contains the eligible training context-plate conditions measured with perturbation *p*. The baseline therefore captures a shared perturbation response without modeling a response specific to the target context.

#### Additive model

The additive model explains an expression response through separate contributions from biological context and perturbation. It allows contexts to differ in their overall response and perturbations to differ in their shared effects, without learning a context-perturbation interaction. Mathematically,

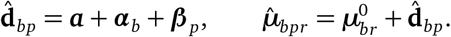

The effects are estimated independently for each gene and sequentially from the training responses. First, ***a*** is the mean response over all training conditions. Next, ***α***_*b*_ is the mean residual over conditions of context *b*. Finally, ***β*** _*p*_ is the mean remaining residual over conditions of perturbation *p*. This sequential procedure defines the effects without additional identifiability constraints.

#### Linear ridge regression

The linear ridge regression baseline represents shared perturbation responses through a low-dimensional description of genes. It constrains the perturbation-mean responses to this gene representation, with regularization shrinking the fitted coefficients.

Following ref. 14, let *D* contain the perturbation-mean responses 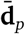 as columns, and let ***ν*** be its row mean. Let *G* be the genes-by-100 matrix of gene scores from the first 100 principal components of the training pseudobulk matrix, whose columns contain mean raw-count profiles centred within each condition. Let *P* be the identity matrix encoding perturbations as one-hot vectors.

We approximate

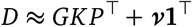

using the regularized closed-form estimator of the reference implementation,

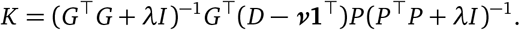

We fix λ = 0.1 to the reference implementation’s value rather than selecting it on validation data. No gradient-based optimization is required.

Writing **e**_*p*_ for the one-hot vector selecting the relevant response column, the prediction is

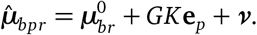

The fitted perturbation response is shared across eligible biological contexts.

#### Nearest neighbor

We introduce a nearest-neighbor baseline that predicts a perturbation’s effect by copying its observed cells from a biological context with similar responses to other perturbations. For each training condition, we compute a response by subtracting the matched control mean from the perturbed mean after applying log(1 + *y*) to individual cell counts. Responses measured on multiple plates are averaged in proportion to their perturbed cell counts, then projected onto 100 principal axes fitted to the training responses. We compare two contexts using their average cosine distance over shared training perturbations with nonzero projected responses. To predict a held-out pair (*b, p*), we select the closest eligible training context measured with *p*. Candidates must share at least one usable training perturbation with the target context. This procedure applies to both biological-context and perturbation generalization.

### Hierarchical balanced stratified sampling

To construct reduced training sets with fewer experimental conditions while maintaining approximately balanced representation of the biological contexts and perturbations axes, as in Figure 3, we use a hierarchical balanced stratified sampling (HBSS) procedure. Unlike uniform sampling, which preserves imbalances in the original dataset, HBSS assigns higher selection probabilities to underrepresented combinations. For example, in the Tahoe-100M biological-context split, the five evaluation cell lines have substantially fewer training combinations than the other cell lines because most of their perturbation responses were withheld. Uniform sampling would preserve this imbalance, whereas HBSS increases their representation in the subsampled training set. Only perturbed combinations are sampled; all control cells remain in training.

#### Inclusion weights

Let *Ω* be the *N* measured combinations (*b, p*, ℓ) of cell line *b*, compound *p* and dose ℓ, with plates pooled, and let *n* = round(*f N*). Starting from *Q*_*bp*ℓ_ = *n*/*N* on *Ω* and 0 elsewhere, iterative proportional fitting rescales *Q* along each axis in turn towards uniform sums (*n*/|*B*| per line, *n*/|*P*| per compound, *n*/|*L*| per dose), clipping every entry to at most 1 after each rescaling so that *Q*_*bp*ℓ_ remains an inclusion probability. Iteration stops once no entry changes by more than 10^−4^ in a sweep.

#### Quotas

Each compound-dose pair receives an integer number of cell lines *q*_*p*ℓ_ by largest-remainder rounding of Σ_*b*_ *Q*_*bp*ℓ_, so that Σ*q*_*p*ℓ_ = *n*; a quota exceeding the number of cell lines measured with the pair is capped there and the excess passed to pairs with unused cell lines.

#### Selection

Within each pair, cell lines are drawn one at a time without replacement with probability proportional to max{*Q*_*bp*ℓ_ − λ*c*_φ(*b*)_, 0}, where *c*_φ(*b*)_ counts the lines already drawn from the tissue type of *b* for this pair and λ = 0.5; if every remaining score is zero, the next line is drawn uniformly. Because λ exceeds most weights, a second cell line from a tissue type is drawn only once every other tissue type has one, with a goal of increasing tissue type.

#### Settings

Tissue types are defined by the cell lines’ organ annotations. The 5% and 25% selections were drawn independently (seed 0). In the PBMC atlas, every donor-cytokine combination is measured, and training combinations are uneven only because held-out donors contribute fewer cytokines; HBSS there balances donors and cytokines without a tissue term.

### Biological program recovery with Hallmark GSEA

To assess whether GENESIS recovers biological programs under an alternative gene-set collection and scoring method, we complemented the PROGENy analysis with preranked GSEA ^42^ of the 50 MSigDB Hallmark gene sets (v2026.1^43^). GENESIS was trained on 15% of training conditions using a 5,750-gene panel covering 4,376 of 4,384 Hallmark genes; additive and nearest-neighbor baselines used the same training conditions. Evaluation covered 75 compounds at 5 µM across five held-out cell lines (375 conditions), each measured on at least two plates with at least 50 perturbed and 50 control cells per plate. GENESIS outperformed both baselines across gene sets, compounds and cell lines. Across gene sets, its mean Pearson correlation with measured responses was 0.32, approaching experimental reproducibility of 0.38 (Extended Data Fig. 13).

Plate-specific responses followed the pseudobulk and library-depth procedures in Methods. Genes were retained at a control detection fraction of at least 10%, averaged equally across plates. Predictions used the measured detection filter and depth adjustment, with additional source-line detection filtering for nearest neighbor. Genes were ranked by adjusted log-fold change, resolving ties by gene name, and scored using preranked GSEA (GSEApy ^44^), with all 50 gene sets retained. Normalized enrichment scores were evaluated using Pearson correlations across gene sets, compounds and cell lines, following the cross-plate comparisons and cluster-bootstrap procedures described in Methods. For the compound measured on three plates, statistics were averaged over three complete analyses, each using a different plate pair.

### Ablations

#### Conditioning covariates in Tahoe-100M

GENESIS generates predictions conditional on all covariates available. To assess the contribution of different conditioning covariates, we trained four GENESIS models using progressively complete inputs: plate alone; plate and cell line; plate, compound and dose; or all four covariates. All four models were trained on the faster configuration (generated repeats per true cell *m* = 96) and were compared only with one another (Extended Data Fig. 15).

#### Energy score versus Gaussian-kernel MMD

To test the effect of the training objective, we retrained GENESIS on the 25% training-cell subset used in Fig. 3a, replacing the energy score and variogram with a Gaussian-kernel MMD objective. MMD is commonly used to fit cell distributions ^3,4^. With State’s energy-distance kernel, MMD^2^ equals energy distance ^3^; we therefore used a Gaussian kernel to compare distinct objectives. The replacement loss was

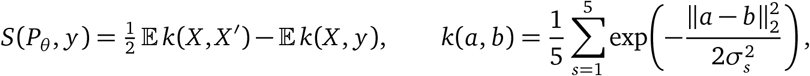

where *X, X* ^′^ are independent model samples and 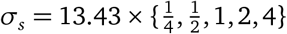. The base bandwidth was fixed before training as the median pairwise distance among 4,096 training cells in log(1+count) space. Averaged over observed cells, this strictly proper score equals 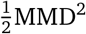 up to a constant.

Each training step used seven conditions, with up to 256 observed and 256 generated cells per condition, weighted by each condition’s training-cell count. We trained for 280,000 optimizer steps, compared with 278,120 for GENESIS. This condition-level objective used up to 1,792 generated cells per step; GENESIS’s per-observed-cell objective used 458,752. Alongside the standard metrics, we evaluated both models using unbiased MMD^2^ with a single Gaussian kernel, setting its bandwidth to the median pairwise distance in each comparison (Extended Data Fig. 16).

### Effect of training settings

We assessed the performance cost of reducing the number of generated cells and the network size, and examined the contribution of the variogram term. Starting from the model of Fig. 2, we varied each setting separately while keeping the remaining training settings fixed.

#### Number of generated cells

We evaluated whether fewer generated cells per observation could reduce training cost while retaining predictive performance. We tested *m* ∈ {16, 64, 256}, with *m* = 256 used in the main model. At the fixed minibatch size of 1,792 observed cells, these settings generate 28,672, 114,688 and 458,752 cells per step, respectively. All generated cells contributed to the energy score and variogram estimators. Five-epoch training times were approximately 4.3, 11.0 and 43.3 hours, respectively (Extended Data Fig. 14a).

#### Variogram weight

We introduced the variogram term to emphasize gene-gene dependencies during training. To assess its contribution, we evaluated weights λ ∈ {0, 0.1, 0.3, 0.5, 1}, including the energy-score-only objective at λ = 0 and the main model’s setting of λ = 0.5 (Extended Data Fig. 14b).

#### Network architecture

We evaluated whether smaller networks could retain the performance of the main architecture. We compared zero hidden layers, one hidden layer (8,192 units) and the main model’s two hidden layers (4,096 and 8,192 units), comprising 2.7, 27.0 and 55.3 million parameters, respectively. All networks received a 1,280-dimensional input comprising four 256-dimensional embeddings (cell line, compound, dose and plate) and a 256-dimensional noise vector. The output head was a linear map to 2,000 genes followed by ReLU, acting directly on the input in the zero-hidden-layer model. Five-epoch training times were approximately 16.8, 31.3 and 43.3 hours, respectively (Extended Data Fig. 14c).

## Extended Data

**Extended Data Fig. 1.**
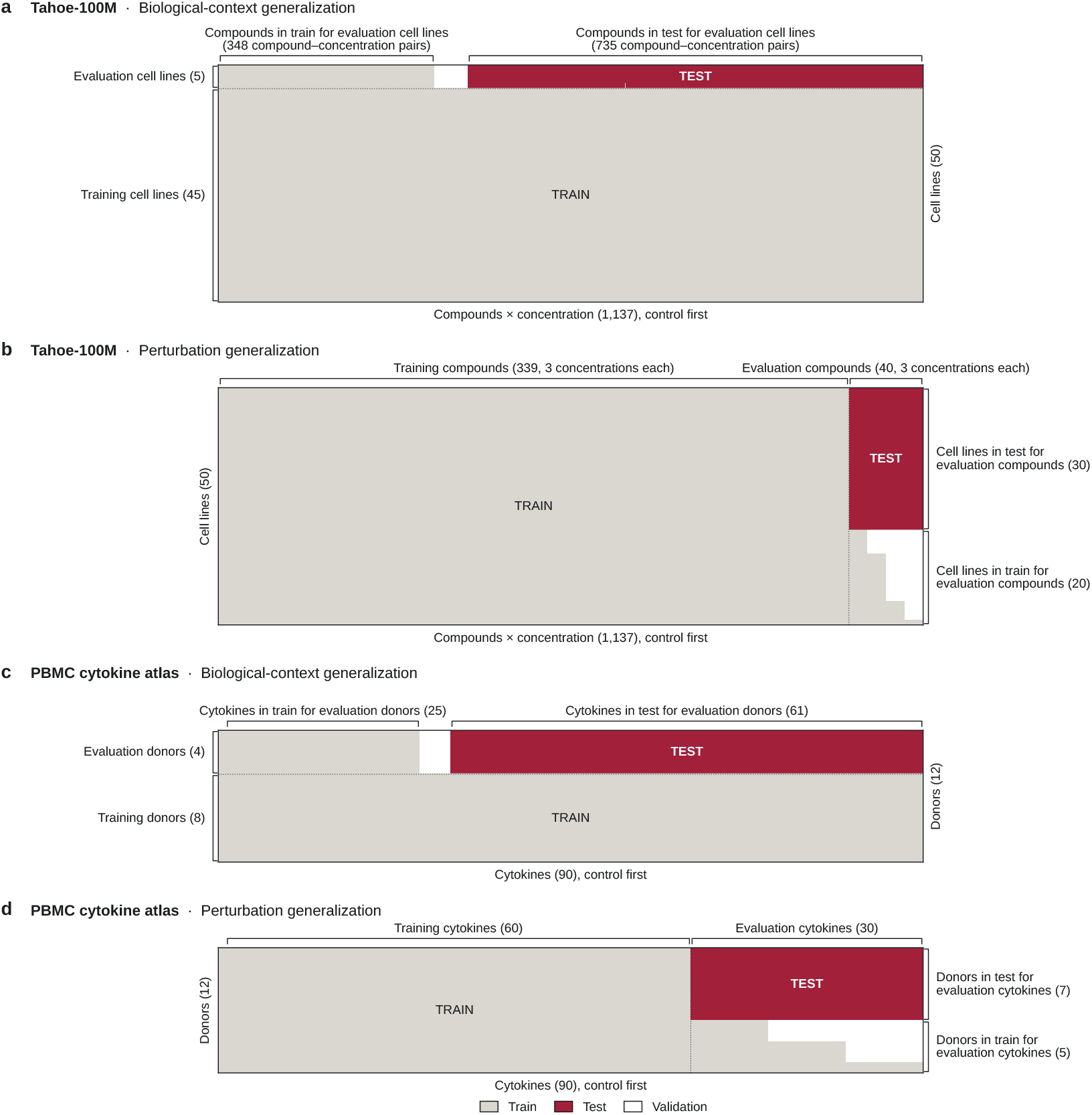
Illustration of training and test splits for biological-context and perturbation generalization. Matrices show training (light grey), test (crimson), validation (white) combinations of biological contexts and perturbations. Columns represent compound–dose pairs in Tahoe-100M and cytokines in the PBMC atlas; controls appear first and remain in training for all contexts. **a**,**c**, Biological-context generalization. Five evaluation cell lines in Tahoe-100M (**a**) and four evaluation donors in PBMC (**c**) retain a minority of perturbation responses for training, with the majority reserved for testing. The remaining 45 cell lines and eight donors provide training responses across the perturbation panel. **b**,**d**, Perturbation generalization. Forty evaluation compounds, at all three doses, are withheld in 30 Tahoe-100M cell lines (**b**); each compound is observed during training in 1, 5, 15 or 20 of the remaining cell lines. Thirty evaluation cytokines are withheld in seven PBMC donors (**d**); each is observed during training in 1, 3 or 5 of the remaining donors. Unused combinations are assigned to validation, and all other perturbations remain in training across contexts. Below the test blocks in **b**,**d**, assignments are sorted independently within each column to show training coverage; rows in these regions therefore do not represent fixed cell lines or donors.

**Extended Data Fig. 2.**
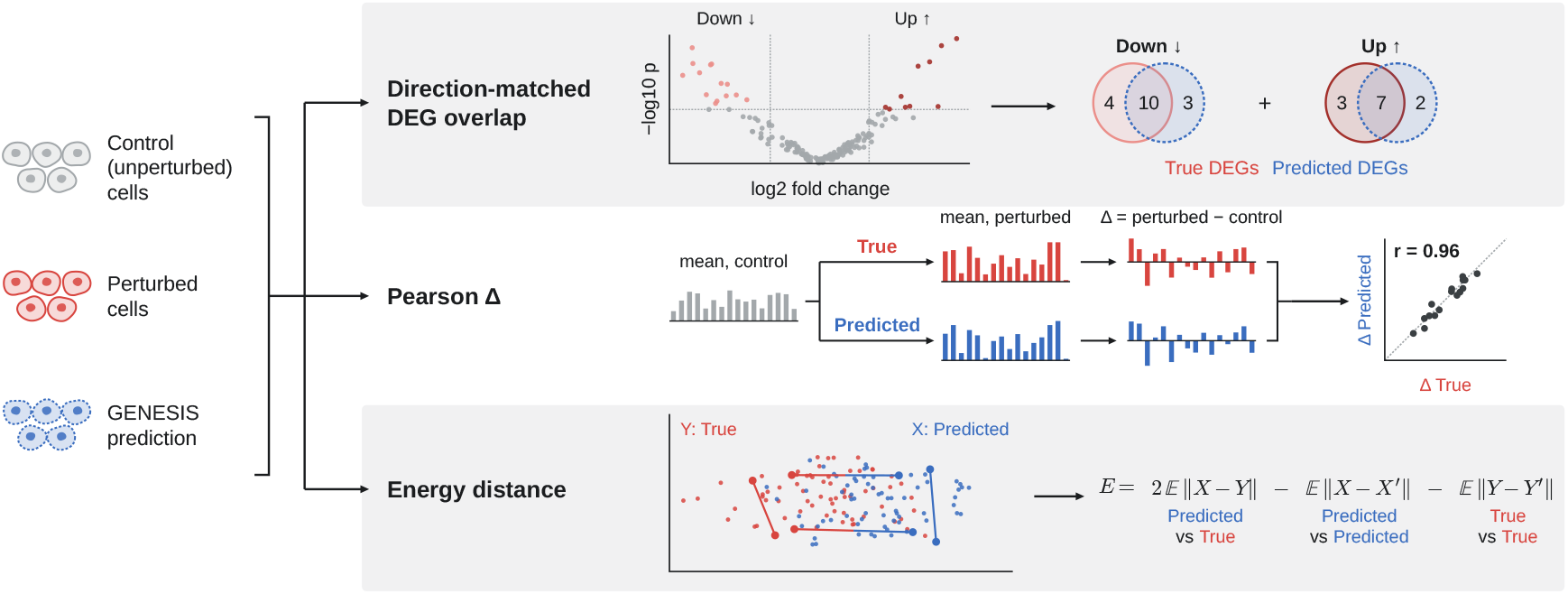
Metrics for evaluating predicted perturbation responses. Observed perturbed cells (red), GENESIS-generated cells (blue) and unperturbed controls (grey) illustrate three metrics. **DEG overlap** measures how well the prediction recovers the most significant differentially expressed genes, requiring agreement in the direction of change. **Control-relative Pearson correlation** measures agreement between observed and predicted mean expression shifts relative to controls. Both metrics use separate, randomly selected halves of the matched controls for the observed and generated populations. **Energy distance** compares the full observed and generated cell distributions using distances between individual cells, without averaging cells or using controls. Higher DEG overlap and Pearson correlation, and lower energy distance, indicate better predictions.

**Extended Data Fig. 3.**
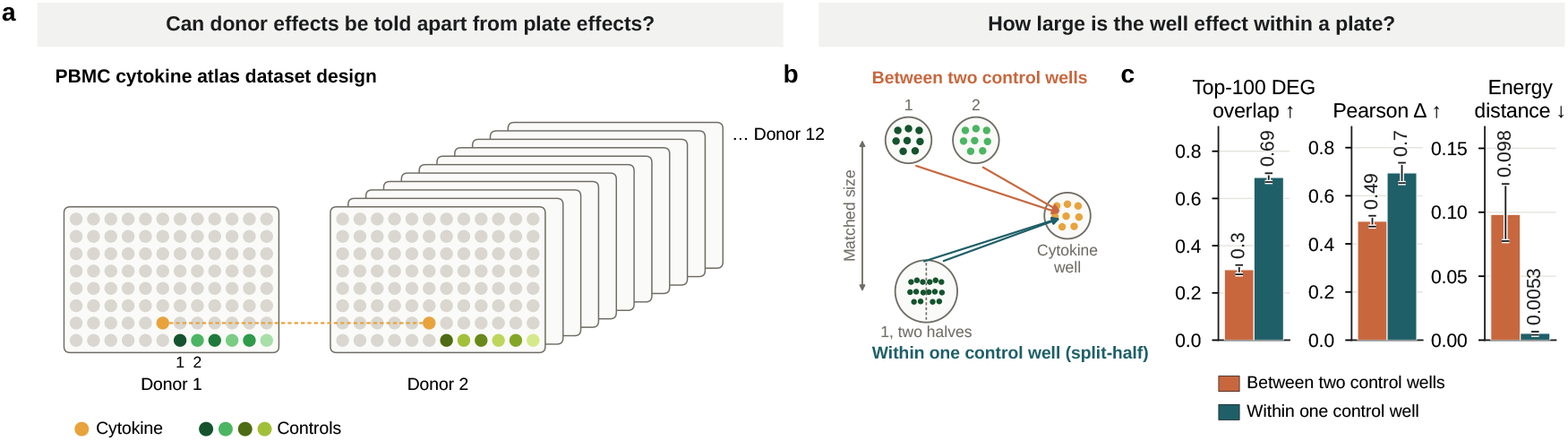
Donor–plate confounding and control-well variability in the PBMC cytokine atlas. **a**, Dataset design. Each of the 12 donors is measured on a separate plate, confounding donor and plate effects. Each cytokine occupies the same well position across plates (orange; dashed line) and is measured once per donor, without replication. Six PBS control wells occupy the edge row of each plate (green). **b**, Comparison of between-well and within-well control variability. The same cytokine-treated cells are compared with two equal-sized control samples drawn either from different control wells (orange) or from disjoint halves of the same well (teal). Agreement between the resulting DEG lists and control-relative mean expression shifts therefore measures sensitivity to the choice of controls. Energy distance is computed directly between the two control samples. **c**, Between-well comparisons yield lower DEG overlap and Pearson *Δ*, and higher energy distance, than within-well comparisons. Analyses include 46 donor-cell-type groups from four donors and 2,805 cytokine-treated conditions (2,792 for DEG overlap). Within each group, all usable control-well pairs and individual wells are sampled 10 times, matching control-sample sizes between comparisons. DEG overlap and Pearson *Δ* are averaged over subsamples for each condition; energy distance is averaged within each group. Bars show means; whiskers indicate 95% confidence intervals from bootstrapping donor-cell-type groups. Between-to within-well mean ratios are 0.43 for DEG overlap (*P* = 2 × 10^−31^), 0.71 for Pearson *Δ* (*P* = 9 × 10^−24^) and 18 for energy distance (*P* = 7 × 10^−22^), using two-sided tests clustered by donor and cell type.

**Extended Data Fig. 4.**
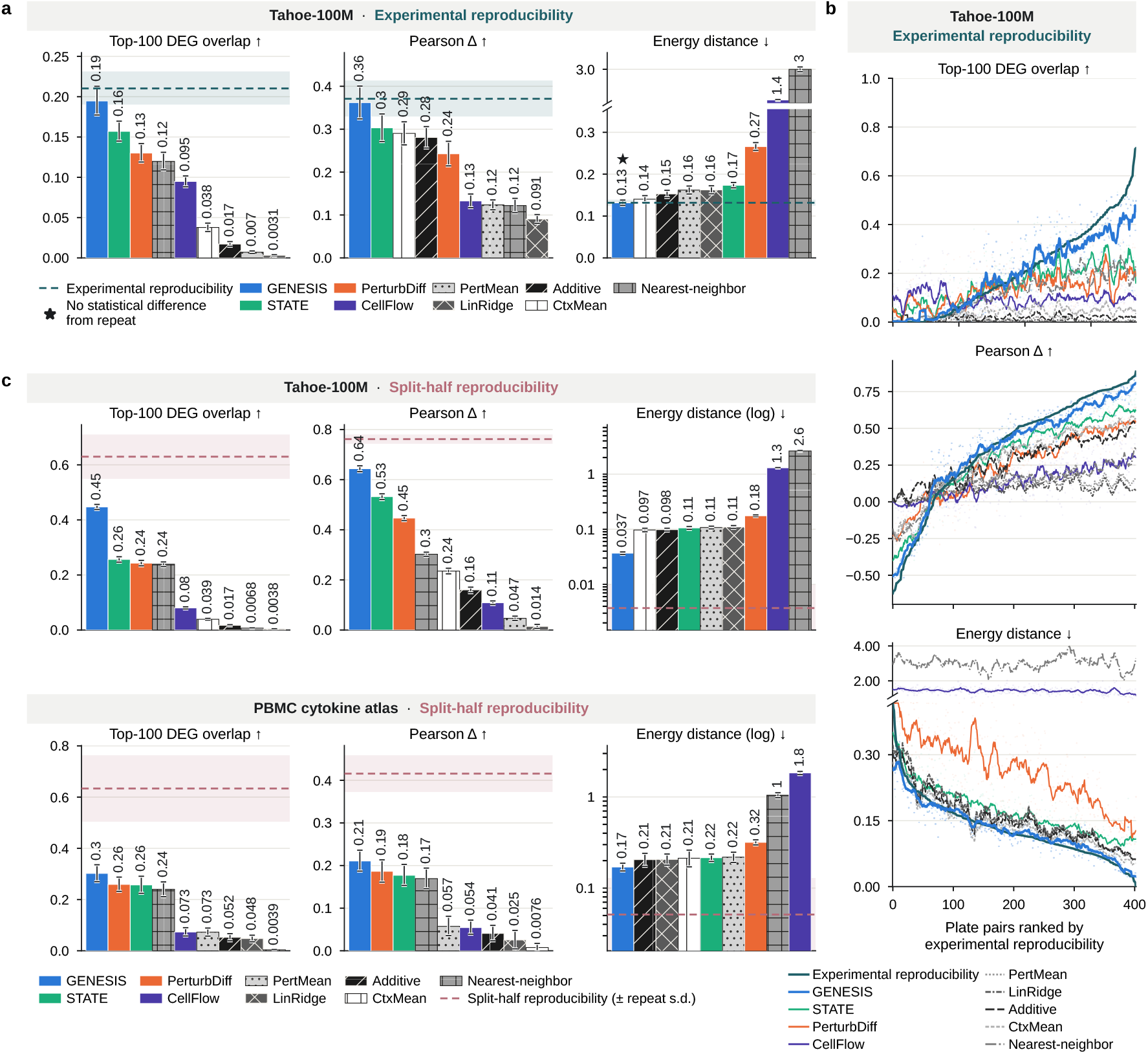
GENESIS achieves state-of-the-art performance approaching experimental reproducibility in the perturbation-generalization setting. Panels follow Fig. 2 except reflecting the perturbation-generalization setting rather than the biological-context generalization split. **a**, Agreement with the experimental repeat on the Tahoe-100M held-out combinations measured on two plates with at least 50 perturbed cells each (403 plate pairs): each method generates each plate from that plate’s control cells and is scored against the other plate. Each metric uses the pairs on which the repeat and every method define it, so DEG overlap, undefined where the repeat has no significant genes, covers fewer pairs (369). Bars, mean of each method; dashed line, the experimental repeat; ↑/↓, higher/lower is better. Above each panel, GENESIS’s ratio of means to the repeat (*R*) and its win rate against it, with two-sided, combination-clustered *P*; n.s., no difference between a method and the repeat detected (both *P* ≥ 0.05). **b**, The same plate pairs ranked by the agreement between experimental repeats; lines, running means; dots, individual pairs. **c**, Against the split-half reference (dashed line; band, mean per-condition s.d. over 20 partitions) on the full test sets: Tahoe-100M, 40 compounds in 30 cell lines at three doses (4,023 conditions; top), and the PBMC cytokine atlas, 30 cytokines in 7 donors (3,456 donor-cell-type-well conditions; bottom). n, conditions per metric; DEG overlap is undefined where the observed half has no significant genes, so it covers fewer conditions. Baselines are shown with black-and-white hatching.

**Extended Data Fig. 5.**
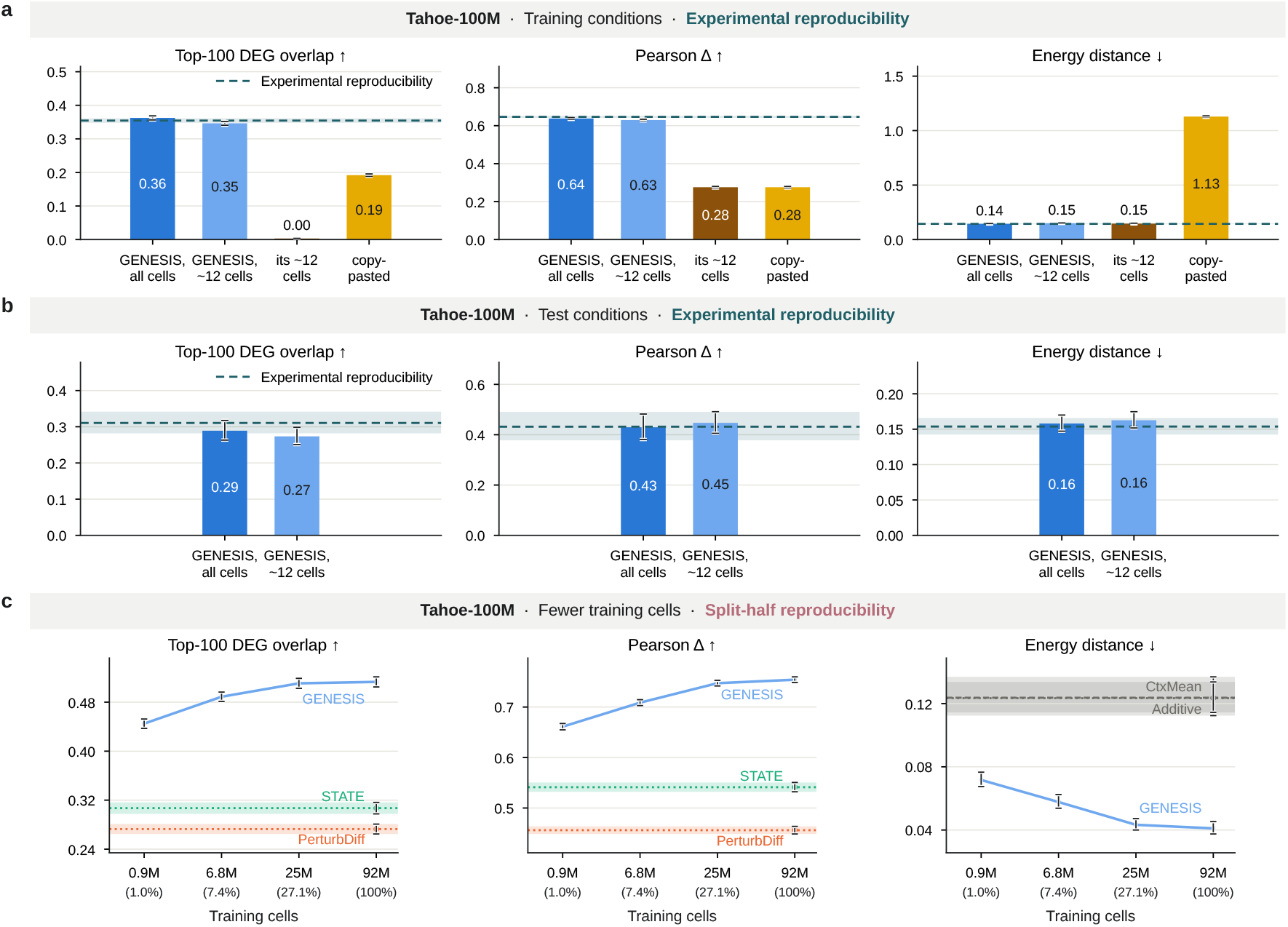
Recovery and generalization from shallow perturbation measurements. **a**,**b**, GENESIS trained on at most 12 perturbed cells per condition, retaining all controls. **a**, Recovery of training-condition responses by generating populations at their original sizes, compared with shallow measurements, replicated shallow measurements and GENESIS trained on all cells. **b**, Generalization to unseen test conditions. Dashed lines in **a**,**b** show experimental reproducibility. **c**, Test performance after subsampling both perturbed and control training cells, evaluated using split-half reproducibility. Dotted lines show the two strongest comparators trained on all data. Points and bars show mean scores; error bars and shaded bands indicate 95% confidence intervals from 10,000 bootstrap resamples of cell line–compound clusters.

**Extended Data Fig. 6.**
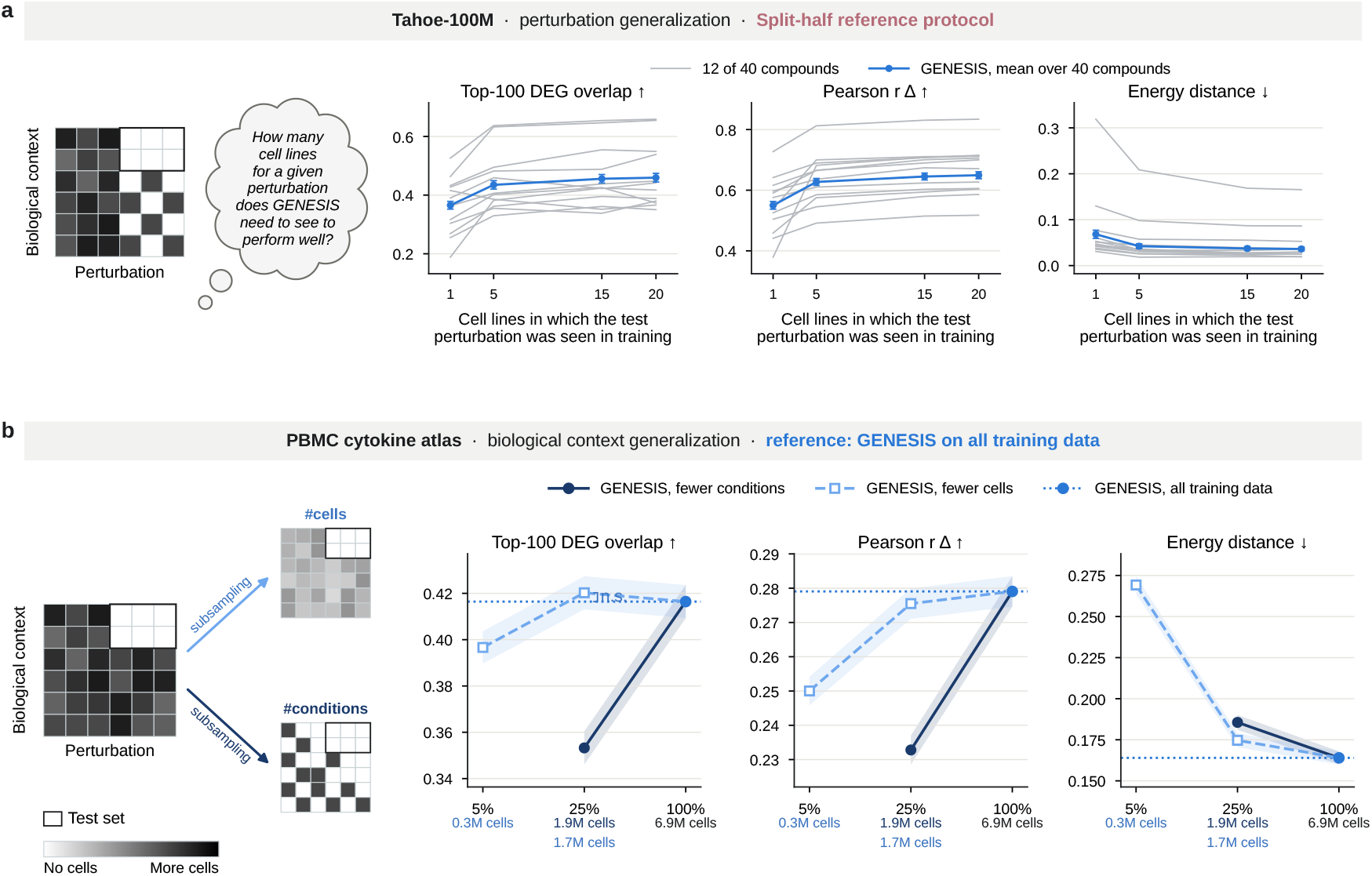
Data efficiency for held-out perturbations and in the PBMC cytokine atlas. **a**, Overlap, as in Fig. 3b but for a held-out perturbation, in Tahoe-100M (perturbation generalization), scored on the full test set by the split-half protocol of Fig. 2c. Each of 40 held-out compounds is observed in training in *k* ∈ {1, 5, 15, 20} of 20 cell lines, rotated over 12 runs so that every compound is scored at every *k*, and is scored at all doses in 30 cell lines in which it is never observed. Gray, 12 individual compounds; blue, mean over compounds ± s.e.m. These runs use the lightweight recipe of Fig. 3b, so absolute values are not those of Extended Data Fig. 4. **b**, Sampling, as in Fig. 3a, in the PBMC cytokine atlas (biological-context generalization): GENESIS trained on 5% or 25% of training cells, or on 25% of donor-cytokine combinations, against GENESIS trained on all the data (dotted line and blue point), scored on held-out donors. The atlas has no experimental repeat, so the full model is the reference. Tick labels, the fraction of training data kept and the training cells (at 25%, fewer conditions above fewer cells); bands, ± s.e.m. over conditions.

**Extended Data Fig. 7.**
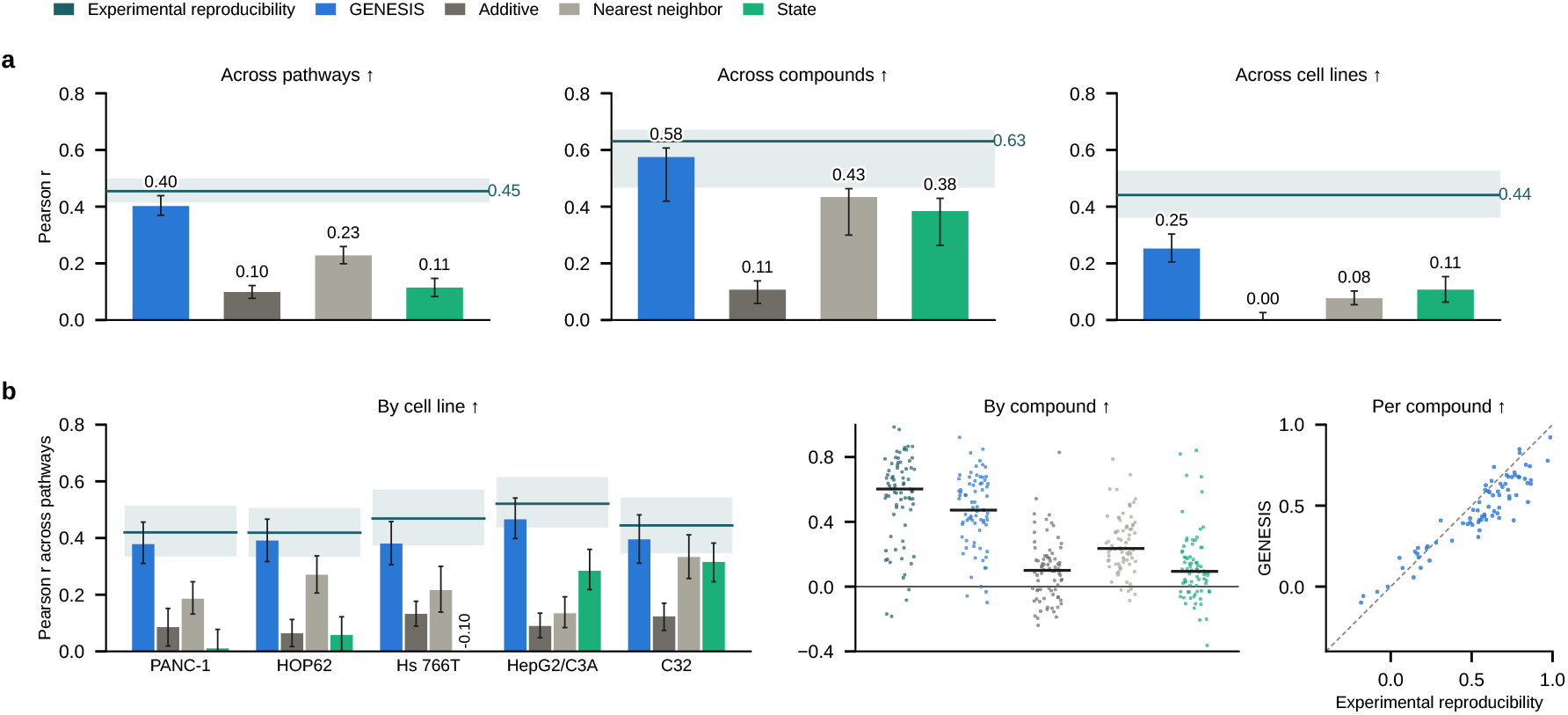
Pathway-level agreement relative to experimental reproducibility. GENESIS and STATE were trained on the same 5% of training cells using the expanded 3,072-gene panel and compared with additive and nearest-neighbor baselines. Responses were scored using the 14 PROGENy pathways and evaluated against experimental reproducibility (Methods). **a**, Mean Pearson correlation across pathways (left), compound–dose conditions (middle), and the five held-out cell lines (right). **b**, Agreement across pathways, broken down by cell line (left) and compound (middle; points show compound-level means across cell lines and doses, with horizontal bars indicating medians). Right, GENESIS agreement versus experimental reproducibility for each compound; the dashed line indicates equality. Teal lines show experimental reproducibility. Error bars and shaded bands indicate percentile 95% confidence intervals from 10,000 cluster-bootstrap resamples, as described in Methods.

**Extended Data Fig. 8.**
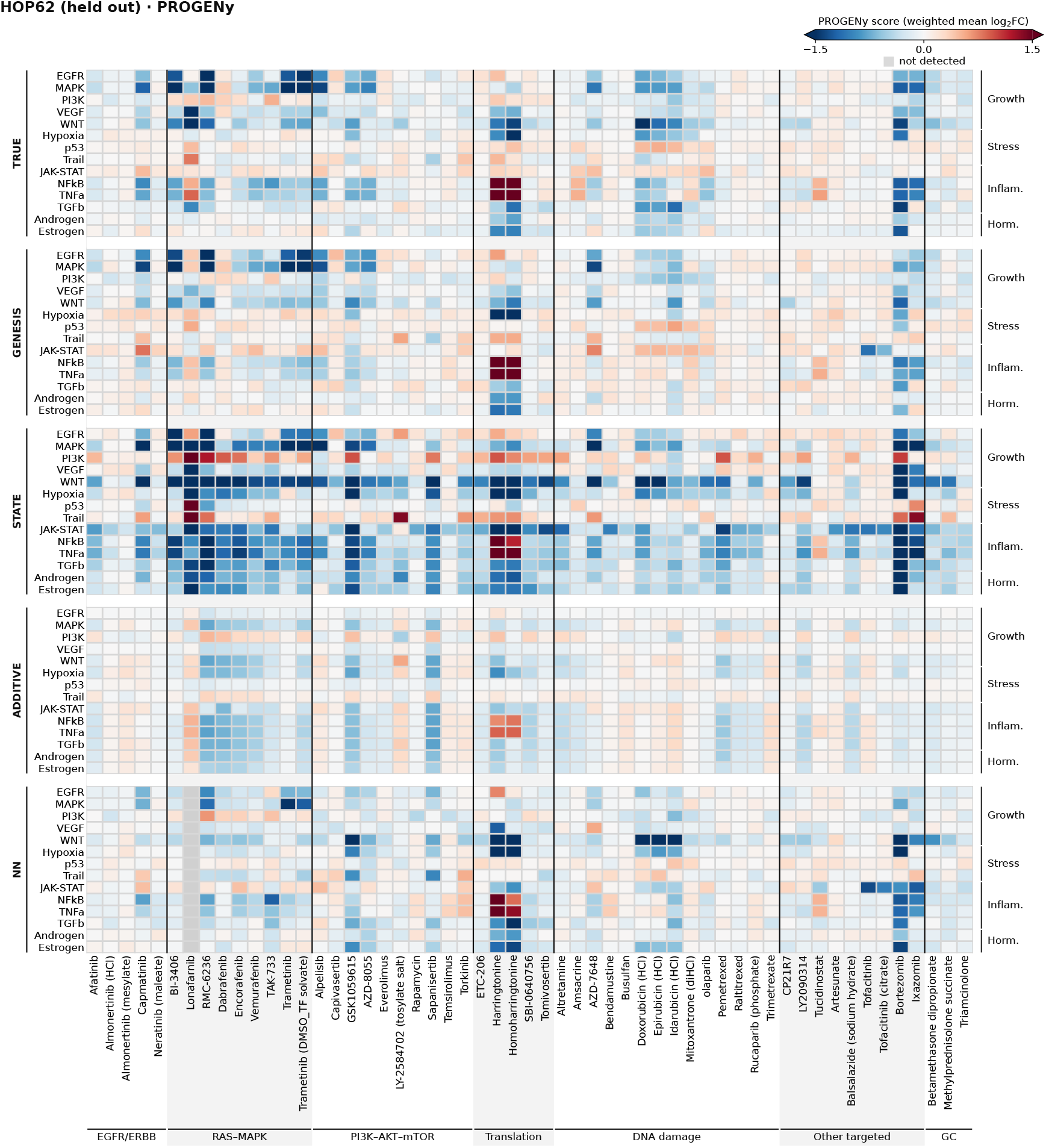
Comparison of predicted pathway responses in HOP62. Measured and predicted PROGENy pathway responses in the held-out HOP62 cell line at 5µM. Heatmaps compare GENESIS, STATE, the additive baseline and nearest-neighbor predictions across 14 pathways and 55 compounds. Colors indicate depth-adjusted, PROGENy-weighted mean log_2_ fold changes relative to matched controls.

**Extended Data Fig. 9.**
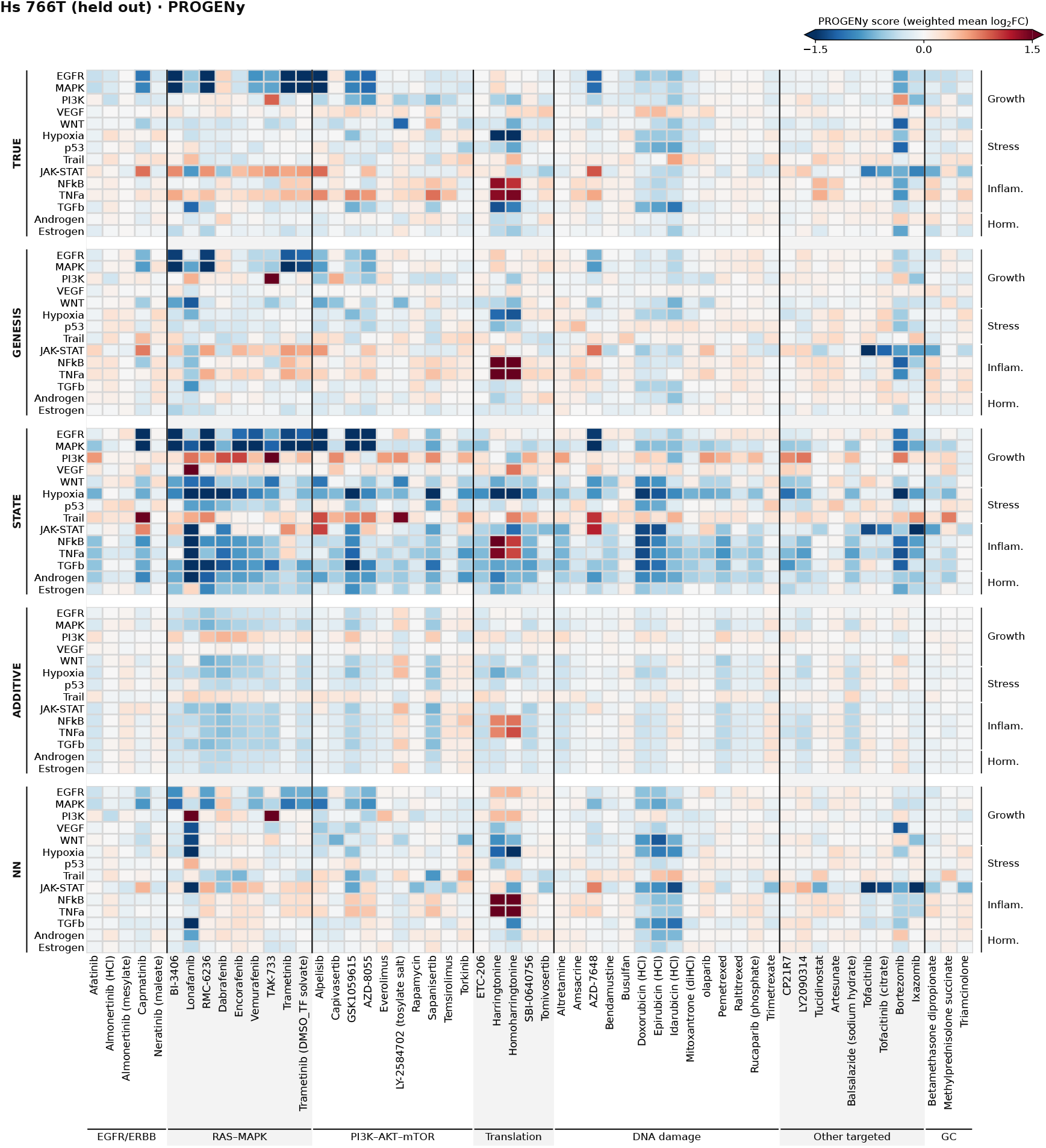
Comparison of predicted pathway responses in Hs 766T. Measured and predicted PROGENy pathway responses in the held-out Hs 766T cell line at 5µM. Models, pathways, compounds and color scale are as in Extended Data Fig. 8.

**Extended Data Fig. 10.**
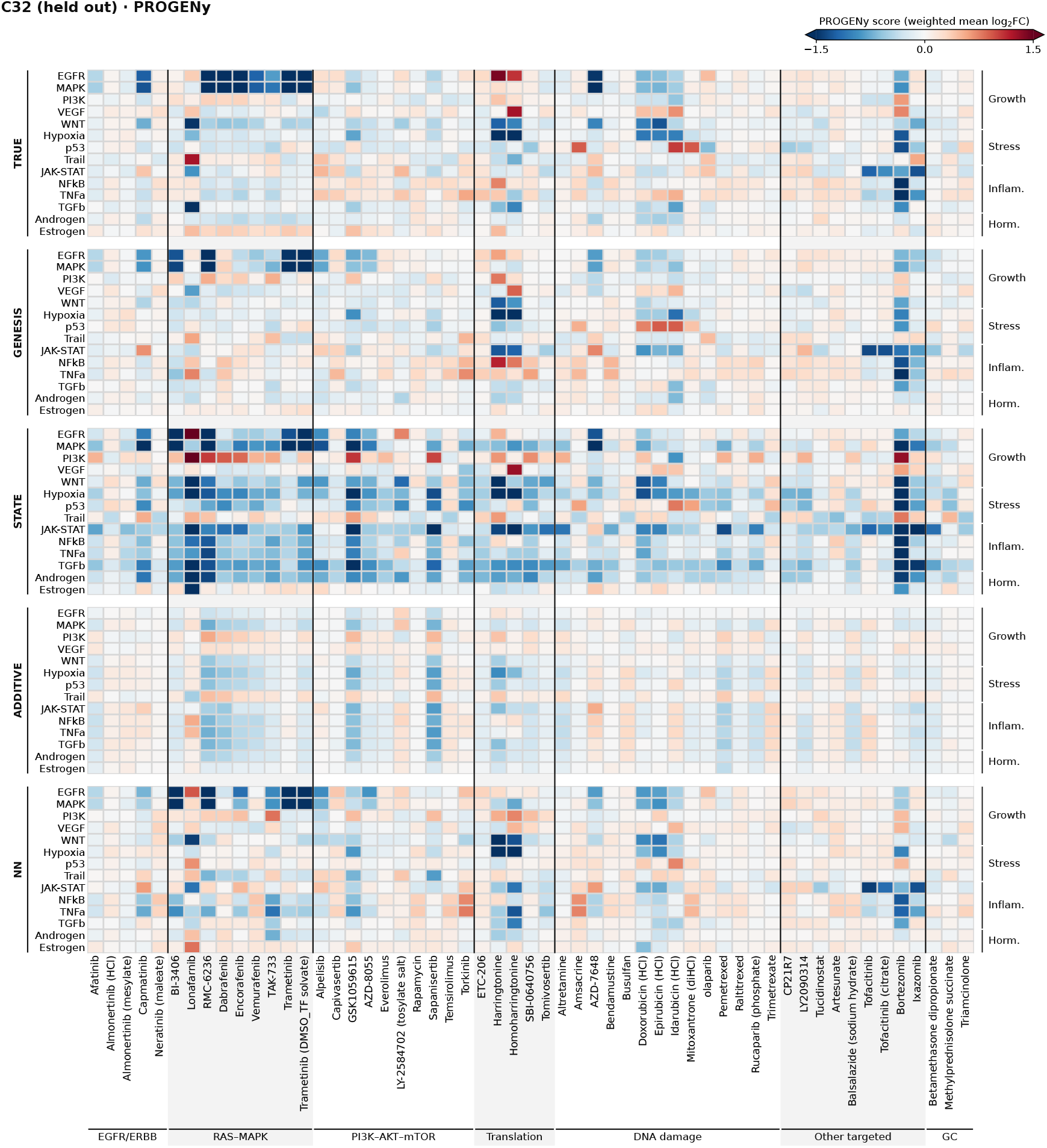
Comparison of predicted pathway responses in C32. Measured and predicted PROGENy pathway responses in the held-out C32 cell line at 5µM. Models, pathways, compounds and color scale are as in Extended Data Fig. 8.

**Extended Data Fig. 11.**
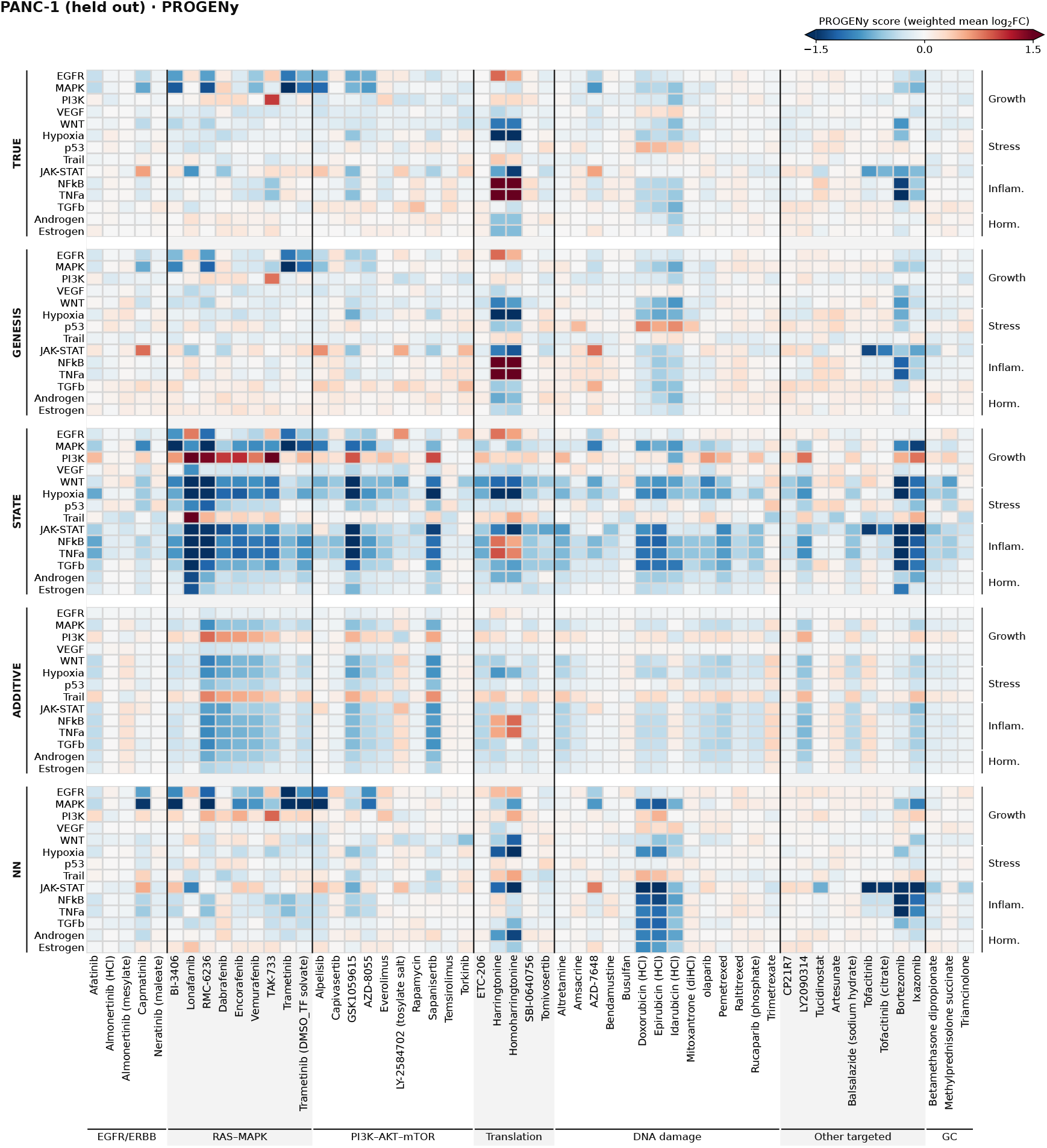
Comparison of predicted pathway responses in PANC-1. Measured and predicted PROGENy pathway responses in the held-out PANC-1 cell line at 5µM. Models, pathways, compounds and color scale are as in Extended Data Fig. 8.

**Extended Data Fig. 12.**
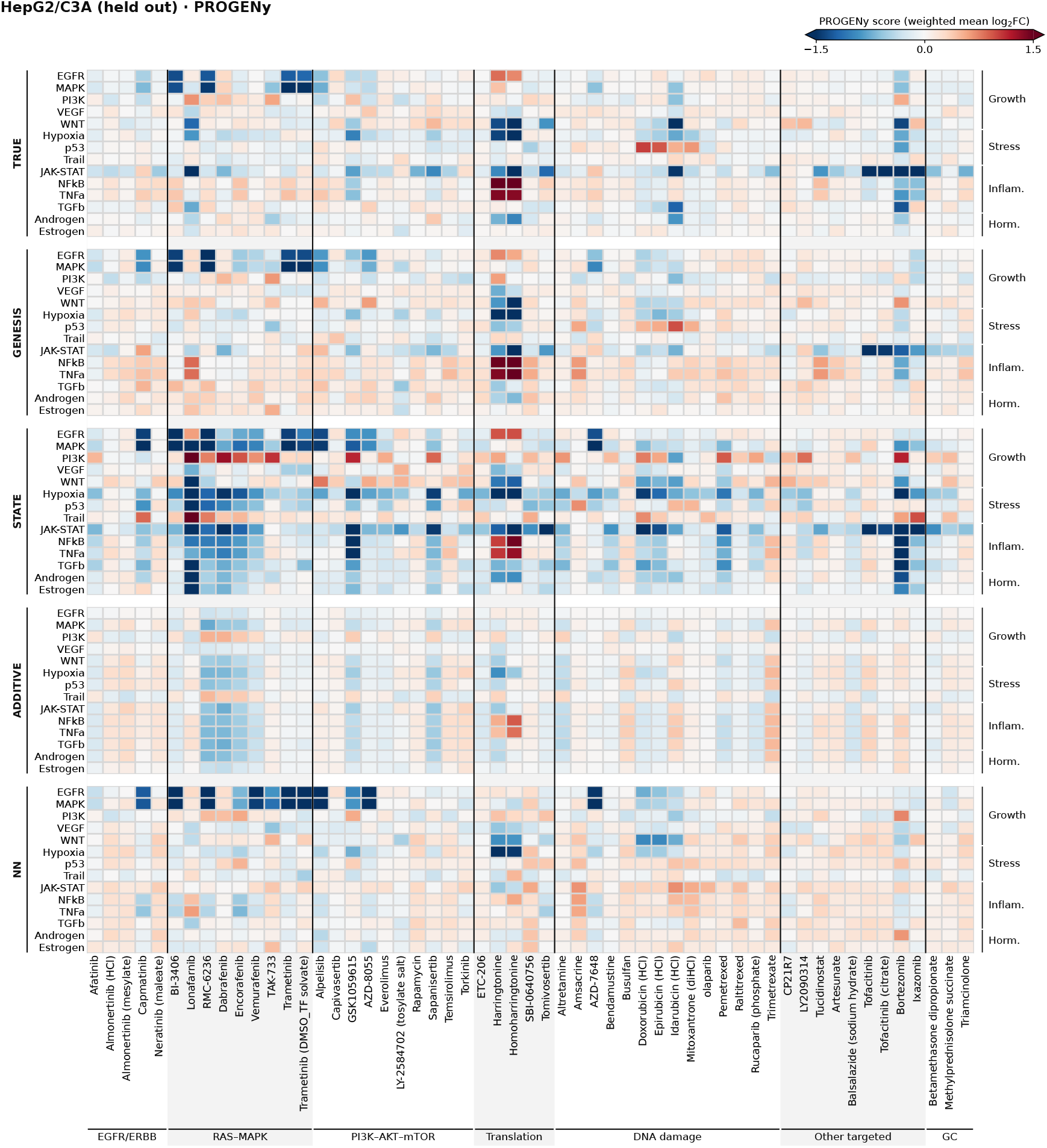
Comparison of predicted pathway responses in HepG2/C3A. Measured and predicted PROGENy pathway responses in the held-out HepG2/C3A cell line at 5µM. Models, pathways, compounds and color scale are as in Extended Data Fig. 8.

**Extended Data Fig. 13.**
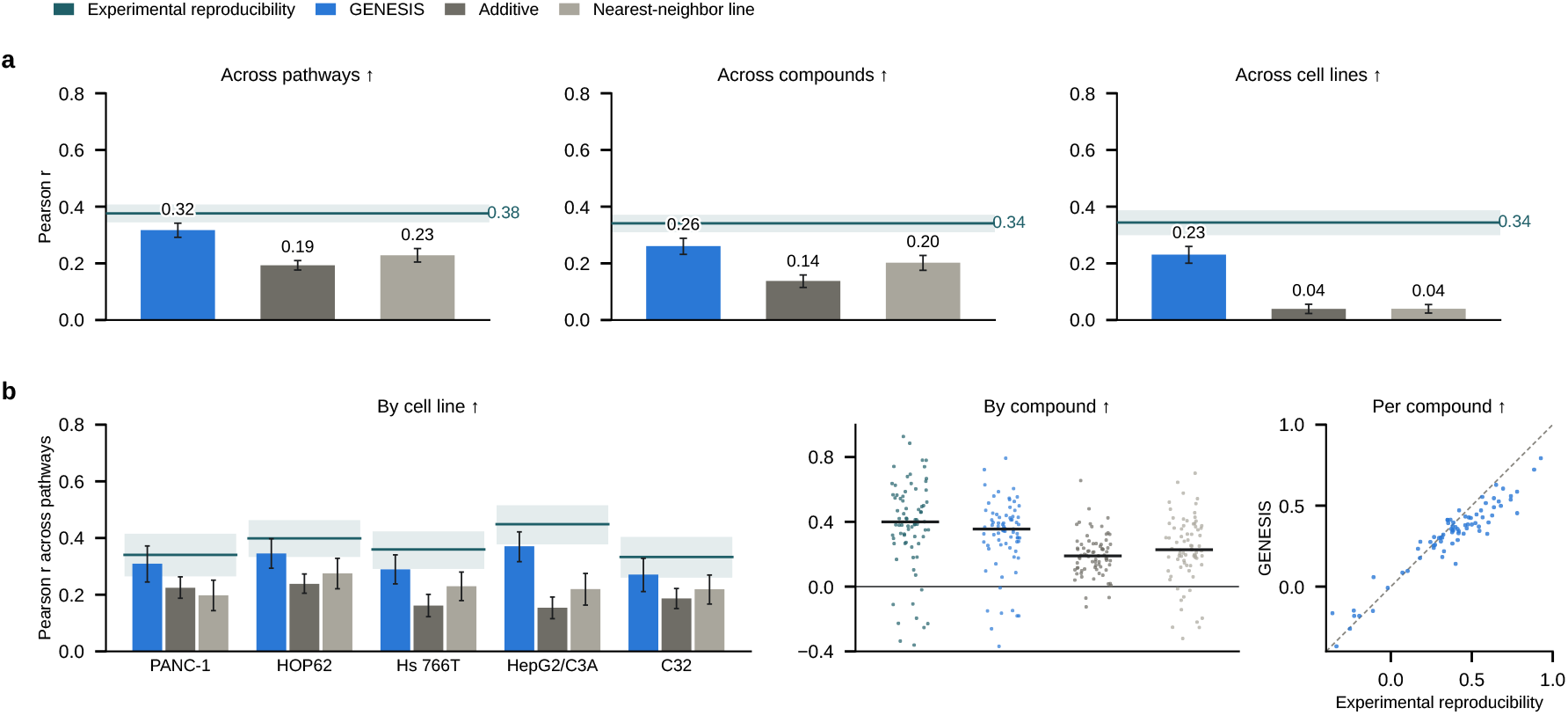
Hallmark pathway agreement across independent plates. Measured and predicted responses were scored by preranked GSEA on 50 MSigDB Hallmark gene sets for 75 compounds at 5 µM across five held-out cell lines (375 conditions; Methods). Predictions were evaluated against the other plate, averaging both directions; teal lines indicate experimental reproducibility. **a**, Mean Pearson (**a**) correlations of normalized enrichment scores across pathways (left), compounds (middle) and cell lines (right). **b**, Across-pathway Pearson correlations summarized by cell line (left), by compound (middle; dots, means across five lines; black bars, medians), and compared between GENESIS and experimental reproducibility for each compound (right; dashed line, identity). Whiskers and shaded bands indicate percentile 95% confidence intervals from 10,000 bootstrap resamples, clustered by cell line-compound for across-pathway and across-compound analyses, and by compound for across-cell-line analyses.

**Extended Data Fig. 14.**
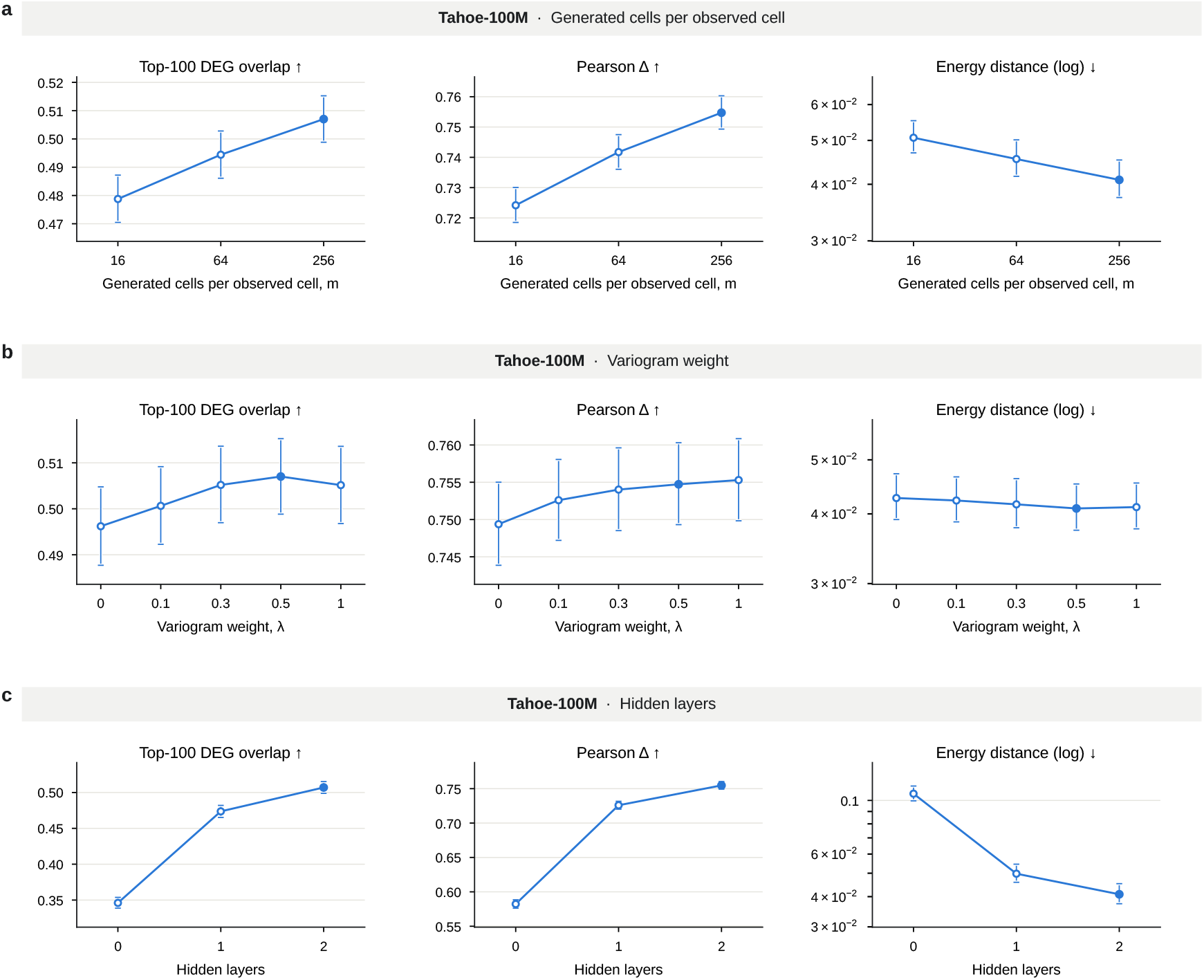
Sensitivity of GENESIS performance to training settings. Models were evaluated on the Tahoe-100M biological-context test split used in Fig. 2c. Each model varied one setting while retaining the remaining settings of the main model. **a**, Number of generated cells per observed cell, *m* ∈ {16, 64, 256}. **b**, Variogram weight, λ ∈ {0, 0.1, 0.3, 0.5, 1}; λ = 0 corresponds to the energy-score-only objective. **c**, Network architecture: zero hidden layers, one hidden layer (8,192 units) or two hidden layers (4,096 and 8,192 units). Filled points indicate the main configuration (*m* = 256, λ = 0.5, two hidden layers) picked during hyperparameter tuning with the validation set.

**Extended Data Fig. 15.**
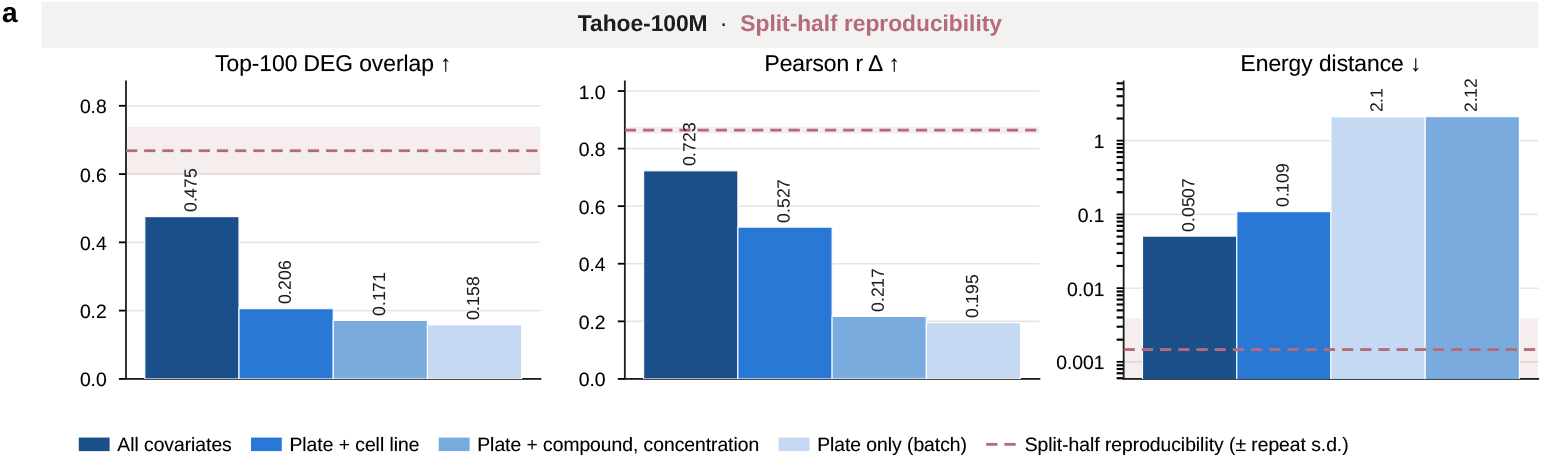
Direct conditioning on experimental covariates improves GENESIS predictions. **a**, Effect of conditioning covariates in Tahoe-100M (biological-generalization split). GENESIS was trained using plate alone, plate and cell line, plate and perturbation (compound and dose), or all covariates. Performance was evaluated on held-out cell lines against the split-half reproducibility reference (dashed line; band, s.d. across partitions).

**Extended Data Fig. 16.**
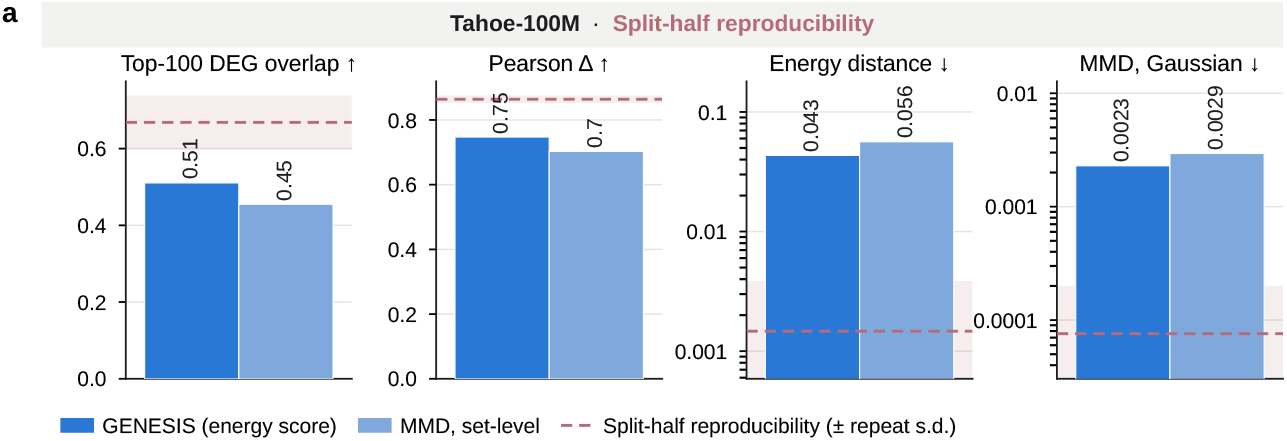
Comparison of the GENESIS and Gaussian-kernel MMD training objectives. GENESIS was trained on the 25% training-cell subset used in Fig. 3a with either its default energy score plus variogram objective or a multiscale Gaussian-kernel MMD objective. Both models used the same architecture; objective estimation and generated-cell budgets are detailed in Methods. Evaluation used the Tahoe-100M biological-context test split and split-half reproducibility as the reference. Bars show mean top-100 DEG overlap, Pearson *Δ*, energy distance and Gaussian-kernel MMD^2^. The evaluation MMD^2^ used a single Gaussian kernel with bandwidth set to the median pairwise distance in each comparison. Bar labels indicate mean scores; dashed lines and shaded bands show split-half reproducibility and its standard deviation across random partitions. Energy distance and MMD^2^ are displayed on logarithmic axes.

## Supplementary Tables

**Supplementary Table 1.**
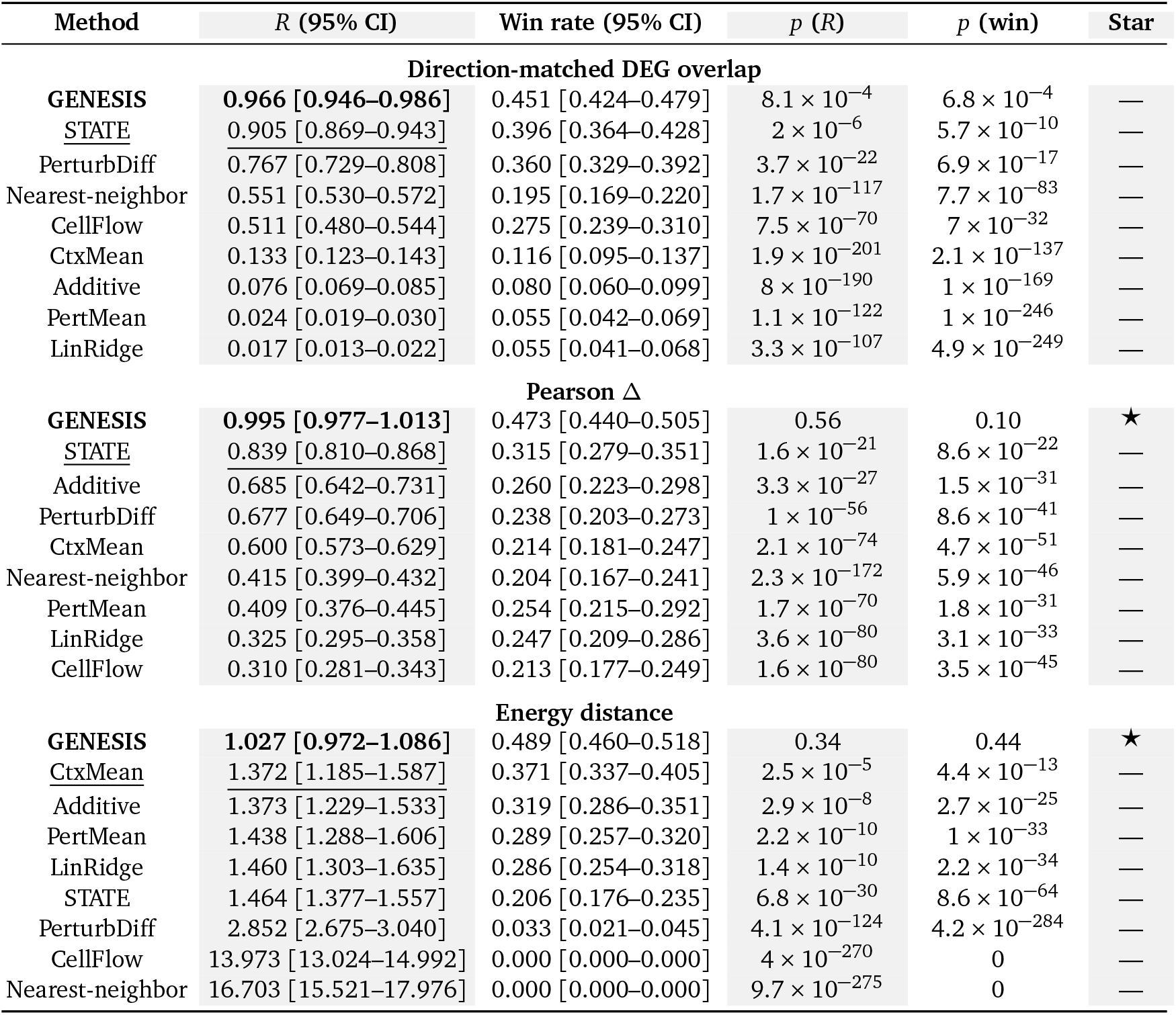
Model performance relative to experimental reproducibility on Tahoe-100M biological-context generalization (Fig. 2a). Predictions were evaluated against observations from the other plate across 530 plate pairs. *R* is the mean model score divided by the mean experimental-repeat score across plates. *R* = 1 indicates equal mean scores; higher values are better for DEG overlap and Pearson *Δ*, and lower values are better for energy distance. Win rate is the fraction of cross-plate predictions that outperform the repeat, counting ties as half. Confidence intervals (95%) and two-sided tests account for clustering by combination, as described in Methods. The tests compare *R* with 1 and win rate with 0.5; stars indicate *p* ≥ 0.05 for both. Methods are ranked by mean performance, with the best in bold and second best underlined.

| Method | *R* (95% CI) | Win rate (95% CI) | *p* (*R*) | *p* (win) | Star |
|---|---|---|---|---|---|
| Direction-matched DEG overlap |  |  |  |  |  |
| **GENESIS** | **0.966 [0.946–0.986]** | 0.451 [0.424–0.479] | 8.1 × 10<sup>−4</sup> | 6.8 × 10<sup>−4</sup> | — |
| STATE | 0.905 [0.869–0.943] | 0.396 [0.364–0.428] | 2 × 10<sup>−6</sup> | 5.7 × 10<sup>−10</sup> | — |
| PerturbDiff | 0.767 [0.729–0.808] | 0.360 [0.329–0.392] | 3.7 × 10<sup>−22</sup> | 6.9 × 10<sup>−17</sup> | — |
| Nearest-neighbor | 0.551 [0.530–0.572] | 0.195 [0.169–0.220] | 1.7 × 10<sup>−117</sup> | 7.7 × 10<sup>−83</sup> | — |
| CellFlow | 0.511 [0.480–0.544] | 0.275 [0.239–0.310] | 7.5 × 10<sup>−70</sup> | 7 × 10<sup>−32</sup> | — |
| CtxMean | 0.133 [0.123–0.143] | 0.116 [0.095–0.137] | 1.9 × 10<sup>−201</sup> | 2.1 × 10<sup>−137</sup> | — |
| Additive | 0.076 [0.069–0.085] | 0.080 [0.060–0.099] | 8 × 10<sup>−190</sup> | 1 × 10<sup>−169</sup> | — |
| PertMean | 0.024 [0.019–0.030] | 0.055 [0.042–0.069] | 1.1 × 10<sup>−122</sup> | 1 × 10<sup>−246</sup> | — |
| LinRidge | 0.017 [0.013–0.022] | 0.055 [0.041–0.068] | 3.3 × 10<sup>−107</sup> | 4.9 × 10<sup>−249</sup> | — |
| Pearson Δ |  |  |  |  |  |
| **GENESIS** | **0.995 [0.977–1.013]** | 0.473 [0.440–0.505] | 0.56 | 0.10 | ★ |
| STATE | 0.839 [0.810–0.868] | 0.315 [0.279–0.351] | 1.6 × 10<sup>−21</sup> | 8.6 × 10<sup>−22</sup> | — |
| Additive | 0.685 [0.642–0.731] | 0.260 [0.223–0.298] | 3.3 × 10<sup>−27</sup> | 1.5 × 10<sup>−31</sup> | — |
| PerturbDiff | 0.677 [0.649–0.706] | 0.238 [0.203–0.273] | 1 × 10<sup>−56</sup> | 8.6 × 10<sup>−41</sup> | — |
| CtxMean | 0.600 [0.573–0.629] | 0.214 [0.181–0.247] | 2.1 × 10<sup>−74</sup> | 4.7 × 10<sup>−51</sup> | — |
| Nearest-neighbor | 0.415 [0.399–0.432] | 0.204 [0.167–0.241] | 2.3 × 10<sup>−172</sup> | 5.9 × 10<sup>−46</sup> | — |
| PertMean | 0.409 [0.376–0.445] | 0.254 [0.215–0.292] | 1.7 × 10<sup>−70</sup> | 1.8 × 10<sup>−31</sup> | — |
| LinRidge | 0.325 [0.295–0.358] | 0.247 [0.209–0.286] | 3.6 × 10<sup>−80</sup> | 3.1 × 10<sup>−33</sup> | — |
| CellFlow | 0.310 [0.281–0.343] | 0.213 [0.177–0.249] | 1.6 × 10<sup>−80</sup> | 3.5 × 10<sup>−45</sup> | — |
| Energy distance |  |  |  |  |  |
| **GENESIS** | **1.027 [0.972–1.086]** | 0.489 [0.460–0.518] | 0.34 | 0.44 | ★ |
| CtxMean | 1.372 [1.185–1.587] | 0.371 [0.337–0.405] | 2.5 × 10<sup>−5</sup> | 4.4 × 10<sup>−13</sup> | — |
| Additive | 1.373 [1.229–1.533] | 0.319 [0.286–0.351] | 2.9 × 10<sup>−8</sup> | 2.7 × 10<sup>−25</sup> | — |
| PertMean | 1.438 [1.288–1.606] | 0.289 [0.257–0.320] | 2.2 × 10<sup>−10</sup> | 1 × 10<sup>−33</sup> | — |
| LinRidge | 1.460 [1.303–1.635] | 0.286 [0.254–0.318] | 1.4 × 10<sup>−10</sup> | 2.2 × 10<sup>−34</sup> | — |
| STATE | 1.464 [1.377–1.557] | 0.206 [0.176–0.235] | 6.8 × 10<sup>−30</sup> | 8.6 × 10<sup>−64</sup> | — |
| PerturbDiff | 2.852 [2.675–3.040] | 0.033 [0.021–0.045] | 4.1 × 10<sup>−124</sup> | 4.2 × 10<sup>−284</sup> | — |
| CellFlow | 13.973 [13.024–14.992] | 0.000 [0.000–0.000] | 4 × 10<sup>−270</sup> | 0 | — |
| Nearest-neighbor | 16.703 [15.521–17.976] | 0.000 [0.000–0.000] | 9.7 × 10<sup>−275</sup> | 0 | — |

**Supplementary Table 2.** GENESIS models trained on reduced data evaluated against experimental reproducibility. Models from Fig. 3a were evaluated across 530 Tahoe-100M plate pairs, scoring predictions against observations from the other plate. Training sets retained fewer perturbed conditions or fewer cells across conditions, with all controls retained. *R* is the ratio of mean model performance to mean experimental-repeat performance across plates; *R* = 1 indicates equal means. Better model performance corresponds to *R* > 1 for DEG overlap and Pearson *Δ*, and *R* < 1 for energy distance. Win rate is the fraction of cross-plate predictions outperforming the repeat, with ties counted as half. We report 95% confidence intervals and two-sided *p* values for *R* = 1 and win rate = 0.5, using the cluster-adjusted tests described in Methods. Stars indicate *p* ≥ 0.05 for both tests.

| Training data | Perturbed cells | *R* (95% CI) | Win rate (95% CI) | *p* (*R*) | *p* (win) | Star |
|---|---|---|---|---|---|---|
| Direction-matched DEG overlap |  |  |  |  |  |  |
| 100% | 89.5M | 0.962 [0.943–0.980] | 0.437 [0.408–0.467] | 8.6 × 10<sup>−5</sup> | 3 × 10<sup>−5</sup> | — |
| 25% conditions | 21.9M | 0.967 [0.948–0.986] | 0.463 [0.434–0.493] | 8 × 10<sup>−4</sup> | 0.01 | — |
| 25% cells | 21.9M | 0.956 [0.937–0.975] | 0.446 [0.417–0.475] | 6.9 × 10<sup>−6</sup> | 2.9 × 10<sup>−4</sup> | — |
| 5% conditions | 4.6M | 0.862 [0.841–0.883] | 0.390 [0.358–0.422] | 1.4 × 10<sup>−29</sup> | 3.8 × 10<sup>−11</sup> | — |
| 5% cells | 4.6M | 0.946 [0.927–0.966] | 0.444 [0.414–0.474] | 2 × 10<sup>−7</sup> | 2.4 × 10<sup>−4</sup> | — |
| Pearson Δ |  |  |  |  |  |  |
| 100% | 89.5M | 0.992 [0.975–1.009] | 0.479 [0.446–0.513] | 0.37 | 0.22 | ★ |
| 25% conditions | 21.9M | 1.004 [0.985–1.023] | 0.481 [0.447–0.515] | 0.71 | 0.27 | ★ |
| 25% cells | 21.9M | 1.000 [0.982–1.019] | 0.475 [0.440–0.509] | 0.99 | 0.14 | ★ |
| 5% conditions | 4.6M | 0.924 [0.902–0.946] | 0.375 [0.339–0.410] | 2.5 × 10<sup>−10</sup> | 9.5 × 10<sup>−12</sup> | — |
| 5% cells | 4.6M | 1.012 [0.989–1.035] | 0.475 [0.439–0.512] | 0.31 | 0.18 | ★ |
| Energy distance |  |  |  |  |  |  |
| 100% | 89.5M | 1.028 [0.973–1.087] | 0.487 [0.459–0.515] | 0.32 | 0.36 | ★ |
| 25% conditions | 21.9M | 1.035 [0.978–1.096] | 0.493 [0.465–0.521] | 0.23 | 0.64 | ★ |
| 25% cells | 21.9M | 1.022 [0.967–1.079] | 0.484 [0.456–0.511] | 0.44 | 0.25 | ★ |
| 5% conditions | 4.6M | 1.152 [1.062–1.250] | 0.402 [0.373–0.431] | 7.1 × 10<sup>−4</sup> | 1 × 10<sup>−10</sup> | — |
| 5% cells | 4.6M | 1.033 [0.973–1.096] | 0.472 [0.442–0.501] | 0.29 | 0.06 | ★ |

**Supplementary Table 3.**
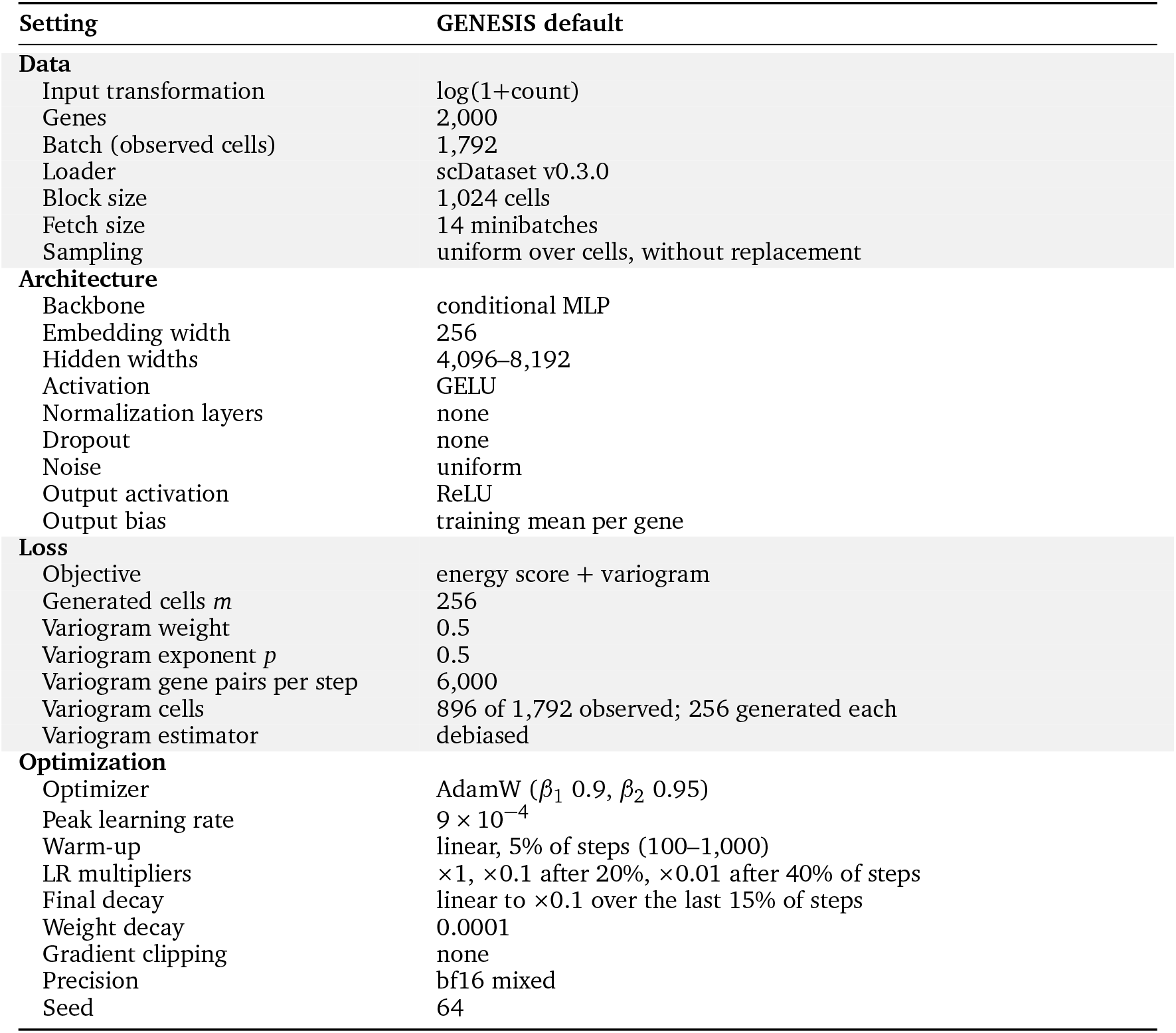
Default GENESIS training settings, as used for the Tahoe-100M benchmark model (Fig. 2). Analysis-specific changes and training durations are described in Methods.

| **Setting** | **GENESIS default** |
|---|---|
| **Data** |  |
| Input transformation | log(1+count) |
| Genes | 2,000 |
| Batch (observed cells) | 1,792 |
| Loader | scDataset v0.3.0 |
| Block size | 1,024 cells |
| Fetch size | 14 minibatches |
| Sampling | uniform over cells, without replacement |
| **Architecture** |  |
| Backbone | conditional MLP |
| Embedding width | 256 |
| Hidden widths | 4,096–8,192 |
| Activation | GELU |
| Normalization layers | none |
| Dropout | none |
| Noise | uniform |
| Output activation | ReLU |
| Output bias | training mean per gene |
| **Loss** |  |
| Objective | energy score + variogram |
| Generated cells *m* | 256 |
| Variogram weight | 0.5 |
| Variogram exponent *p* | 0.5 |
| Variogram gene pairs per step | 6,000 |
| Variogram cells | 896 of 1,792 observed; 256 generated each |
| Variogram estimator | debiased |
| **Optimization** |  |
| Optimizer | AdamW (*β*<sub>1</sub> 0.9, *β*<sub>2</sub> 0.95) |
| Peak learning rate | 9 × 10<sup>−4</sup> |
| Warm-up | linear, 5% of steps (100–1,000) |
| LR multipliers | ×1, ×0.1 after 20%, ×0.01 after 40% of steps |
| Final decay | linear to ×0.1 over the last 15% of steps |
| Weight decay | 0.0001 |
| Gradient clipping | none |
| Precision | bf16 mixed |
| Seed | 64 |

